# The Effect of Pseudoknot Base Pairing on Cotranscriptional Structural Switching of the Fluoride Riboswitch

**DOI:** 10.1101/2023.12.05.570056

**Authors:** Laura M Hertz, Elise N White, Konstantin Kuznedelov, Luyi Cheng, Angela M Yu, Rivaan Kakkaramadam, Konstantin Severinov, Alan Chen, Julius B Lucks

## Abstract

A central question in biology is how RNA sequence changes influence dynamic conformational changes during cotranscriptional folding. Here we investigated this question through the study of transcriptional fluoride riboswitches, non-coding RNAs that sense the fluoride anion through the coordinated folding and rearrangement of a pseudoknotted aptamer domain and a downstream intrinsic terminator expression platform. Using a combination of *E. coli* RNA polymerase *in vitro* transcription and cellular gene expression assays, we characterized the function of mesophilic and thermophilic fluoride riboswitch variants. We showed that only variants containing the mesophilic pseudoknot function at 37 °C. We next systematically varied the pseudoknot sequence and found that a single wobble base pair is critical for function. Characterizing thermophilic variants at 65 °C through *Thermus aquaticus* RNA polymerase *in vitro* transcription showed the importance of this wobble pair for function even at elevated temperatures. Finally, we performed all-atom molecular dynamics simulations which supported the experimental findings, visualized the RNA structure switching process, and provided insight into the important role of magnesium ions. Together these studies provide deeper insights into the role of riboswitch sequence in influencing folding and function that will be important for understanding of RNA-based gene regulation and for synthetic biology applications.

## INTRODUCTION

Cells have evolved a wide array of molecular mechanisms that can sense changes in intra- and inter-cellular chemical signals and respond through regulating gene expression. RNA riboswitches are one such mechanism used by broad ranges of organisms including bacteria, archaea, and some eukaryotes^1^. Riboswitches are RNA-based sensors that regulate gene expression in a ligand-dependent manner through the interworking of two functional domains^2^: an aptamer domain that folds into a structure to form a specific ligand-binding pocket, and a downstream expression platform that can change its conformation based on the aptamer ligand binding state. Aptamer domains that can specifically bind a diverse array of amino acids, nucleotides, cofactors, metals and other compounds exist^1^, and are usually highly conserved within the particular class of ligand they bind^3^. In contrast, expression platforms are inherently more diverse, as different RNA folds can regulate different aspects of gene expression including transcription, translation, and splicing^1,4^. Given the direct relationship between RNA structure and gene regulation, riboswitches have been important model systems used to study and understand RNA structure dynamics, and have found their own specific uses within RNA-based biotechnologies including as antibiotic targets^5^, within artificial cell systems^6^, and as diagnostic biosensors^7^.

While a great deal is known about the RNA structural biology behind aptamer-ligand interactions^8–11^, major questions remain about how ligand binding is converted to RNA structural changes of the expression platform that lead to gene regulation. Especially interesting are transcriptional riboswitches, as aptamer-ligand binding and resulting expression platform structural changes must be relayed on timescales corresponding to transcription elongation rate to appropriately regulate gene synthesis^12^, necessitating a switching mechanism that utilizes cotranscriptional folding^13^. Recent investigations suggest overlapping sequences allow for a strand displacement mechanism^14^, where cotranscriptional rearrangements can occur through the aptamer by the expression platform^15^ or through an intermediate structure competing between the aptamer and expression platform^16^. These have led to observations that nucleotide changes in the expression platform result in altering riboswitch sensitivity and dynamic range^16–19^. Additionally, ligand-binding can dictate mutually exclusive coaxial stacking conformations in the aptamer for a structural orientation mechanism to regulate the riboswitch switching mechanism^20^. However, the influence of changing aptamer sequence, when there is expression platform overlap, on the switching mechanism is still largely unexplored.

Here we sought to investigate questions about the influence of aptamer/expression platform sequences on switching in the context of the transcriptional fluoride riboswitch mechanism. The fluoride riboswitch is a well-characterized system in which the relatively small and compact *crcB* pseudoknot aptamer motif^21^ is stabilized by the coordination of fluoride by three magnesium ions^8,22^ (Figure 1A). Recently, the *Bacillus cereus* (*B. ce*) transcriptional fluoride riboswitch has been used both for biotechnology purpose as a water quality sensor^7^ as well as a model system to study cotranscriptional folding^22–26^. From these cotranscriptional studies, it was revealed that regardless of ligand presence, the aptamer structure forms and it is only through structural rearrangement in the absence of ligand that the expression platform forms an intrinsic termination hairpin^23,24^. Furthermore, it was found that ligand binding leads to anti-termination through the formation of a nested long-range base pair, termed the linchpin, that prevents expression platform folding^24^.

**Figure 1.**
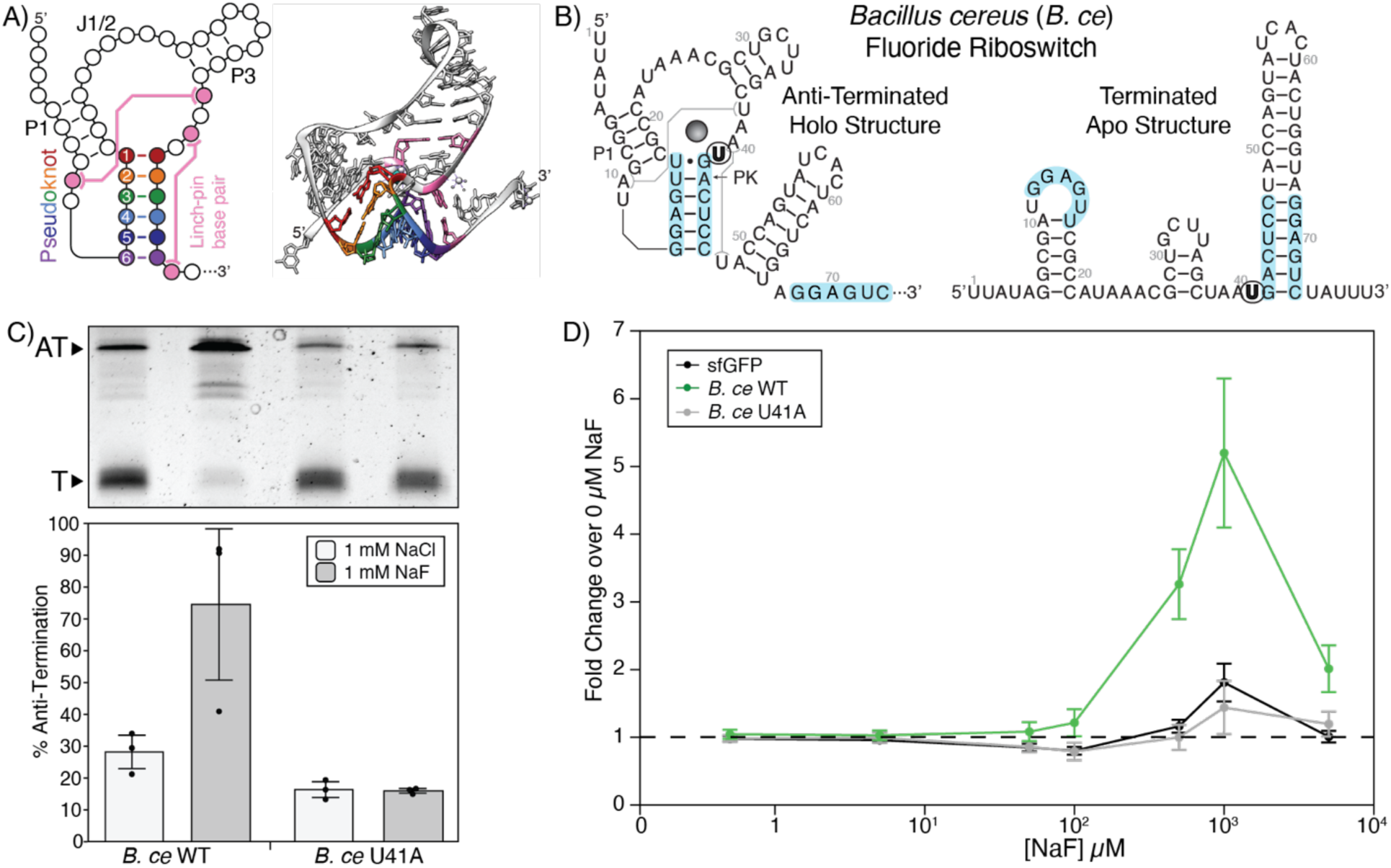
*Bacillus cereus* fluoride riboswitch gene expression assays. A) Secondary and tertiary (PDB ID 4ENC) representation of the fluoride riboswitch aptamer. The pseudoknot is colored in rainbow, with key nucleotide positions given relative numbers to their position in the pseudoknot. Long-range interactions are colored pink. Important structural features are labeled; P1 is the P1 Stem, J1/2 is the region that connects the P1 and P3 stems, and P3 is the P3 Stem. B) Secondary structure of the *Bacillus cereus* wild type fluoride aptamer annotated with sequence (GenBank AE017194.1, bases 4763724 to 4763805^23^). The location of the ligand unresponsive mutation, U41A, is bolded and circled. C) Single-round *in vitro* transcription reactions with *E. coli* RNAP for *B. ce* WT and *B. ce* U41A riboswitches performed with either 1 mM NaCl or 1mM NaF. A representative gel is shown above a plot of band intensity quantification of experimental replicates. The Anti-Terminated (AT) (185 nt) and Terminated (T) (70 nt) bands are indicated. D) Cellular gene expression assay measured with flow cytometry showing a dose response to externally supplied NaF using three constructs: constitutively expressed super-folded GFP (sfGFP), *B. ce* WT fluoride riboswitch, and the ligand unresponsive *B. ce* U41A riboswitch. Fold change was calculated by dividing observed fluorescence at the specific NaF concentration by the observed fluorescence with 0 µM NaF added. A dashed line is drawn at a fold change of 1. Points in C represents three experimental replicates (N=3), with bars representing the average and error bars representing standard deviation. Replicate gels are found in Supplementary Data File 3. Points in D represents the average fold-change of NaF to NaCl for three biological replicates, each analyzed over three technical replicates (N=9) with error bars representing standard deviation. MEFL – molecules of equivalent fluorescein.

While our knowledge of the fluoride riboswitch mechanism is improving, none of the previous studies explored the influence of base pair identity of the critical overlap between the fluoride aptamer pseudoknot and the expression platform (Figure 1B) on fluoride riboswitch function. To address this gap, we therefore sought to investigate the role of sequence variation within the pseudoknot on the switching mechanism of the fluoride riboswitch. Specifically, we combined *in vitro* transcription and cellular gene expression assays to assess how sequence variation within the pseudoknot of the fluoride riboswitch impacted function. The analysis revealed that a wobble base pair at the top of the aptamer is important for conformational switching. We then adapted our *in vitro* transcription assay to study the influence of temperature on structural switching, and found that the wobble base pair continued to be vital for structural switching even at 65 °C. To extend our investigation, we performed all-atom molecular dynamics (MD) simulations to understand how the wobble pair could influence the folding mechanism. The simulations supported the functional data and suggested that the wobble base pair allows relief of the topological twist of the pseudoknot to allow the expression platform to break apart the pseudoknot in the absence of fluoride. We also implemented updated magnesium ion parameters within the simulations to investigate the role of magnesium on the switching mechanism. Analysis of magnesium ions within the simulations revealed that in the presence of fluoride, magnesium appears to concentrate at the structural switching interface, resulting in prevention of the terminator from breaking apart the pseudoknot. Overall, our results add to the growing body of knowledge that individual nucleotides can have drastic impact on RNA folding dynamics and function.

## MATERIALS AND METHODS

### Cloning and plasmid construction

The 79 nt *B. ce* fluoride riboswitch sequence was obtained from GenBank, <u>CP000227.1</u>, bases 4763724 – 4763805 as previously described^23^, while the 82 nt *Thermotoga petrophila (T. pe)* fluoride riboswitch was obtained from GenBank <u>CP000702.1</u>, bases 17948122 to 1794906. Both riboswitch sequences were cloned into plasmid constructs (Supplementary Data File 1) following previous work^7^ by placing them downstream of the consensus *E. coli* sigma 70 promoter sequence (J23119 in the registry of standardized biological parts), and upstream of a translational reporter system comprised of a synthetic ribosome binding site and super-folder GFP^27^ (sfGFP) coding sequence (Supplementary Data File 1). Mutations to the riboswitches and surrounding sequence contexts were constructed through inverse polymerase chain reaction (iPCR). Primers were ordered from Integrated DNA Technologies (IDT). Two hundred µL PCR reactions were prepared with 1 – 10 ng/µL of template plasmid, 250 µM dNTPs, 1X Phusion Buffer, 500 nM of forward and reverse primer, and 0.625 µL of Phusion Polymerase (2,000 U/mL, NEB, #M0530L). The thermal cycling program consisted of an initial 3 min at 98 °C, followed by 10 cycles of 98 °C for 30 sec, a 60-70 °C temperature gradient for 45 sec, and 72 °C for 90 sec, then 28 cycles of 98 °C for 30 sec, a 65 °C step for 45 sec, and 72 °C for 90 sec, finishing off with 72 °C for 5 minutes and holding at 12 °C. PCR reactions were run on a 1% agarose gel to check amplification. Successful reactions were directly incubated at 37 °C with 1 µL of DpnI (NEB, #R0176L) for 30 min to 3 hrs. The reactions were then washed through a DNA column and eluted in 20 µL of water. Seven µL of DNA product was circularized in a reaction with 1 µL of 10X T4 DNA Ligase Reaction Buffer (NEB, #B0202A), 1 µL T4 Ligase (NEB, #M0202L), and 1 µL T4 PNK (NEB, #M0201L) for 30 to 60 min at room temperature. The ligated product was then transformed into NEBTurbo competent *E. coli* cells (NEB, #C2984H) and plated on Lysogeny broth (LB) agar plates with 34 µg/mL chloramphenicol. The plates were incubated overnight at 37 °C. The following day, single colonies were grown in LB media with 34 µg/mL chloramphenicol, mini-prepped, and sequence confirmed using Sanger Sequencing (Quintara).

### DNA template preparation for *in vitro* transcription

DNA templates for single-round *E. coli in vitro* transcription were synthesized through a 200 µL PCR reaction containing 10x Taq Buffer (NEB, #B9014S), 250 µM dNTP Mix (NEB, #N0447L), 125 nM primer A (Supplementary Table S1), 125 nM primer B (Supplementary Table S1), 1 µL DNA plasmid, and 1 µL Taq DNA Polymerase (NEB, #M0273X). The PCR thermal cycling program consisted of an initial 3 min at 95 °C, followed by 25 cycles of 95 °C for 45 sec, 53 °C for 30 sec, and 68 °C for 60 sec, finishing off with 68 °C for 5 minutes and holding at 12 °C. DNA was purified through ethanol precipitation with 20 µL 1 M NaCl, 1 µL glycogen (Thermo, #AM9515), and 600 µL EtOH. The precipitated pellet was air dried, dissolved in H_2_O, and run on a 2% agarose gel. The template band was extracted using a QIAquick Gel Extraction Kit (Qiagen, #28706X4) and re-purified through ethanol precipitation. DNA concentrations were measured on a Qubit 2.0 Fluorometer (Life Technologies, #Q33216).

### Single-round *in vitro* transcription with *E. coli* RNA polymerase

25 µL single-round IVT reactions were assembled with the following concentrations of reagents: 20 mM Tris-HCl pH 8.0, 0.1 mM EDTA pH 8.0, 1 mM DTT, 50 mM KCl, 5 mM MgCl_2_, 0.25 µL BSA, 100 nM DNA template, and 2 µL *E. coli* RNA polymerase holoenzyme (NEB, #M0551S). Either 1 mM of NaCl or NaF was also included depending on the experimental conditions. Once assembled on ice, the reactions were incubated at 37 °C for 10 min to form transcription complexes. Transcription was then initiated with the addition of rNTPs to a final concentration of 500 µM (125 µM of each rNTP) and 0.01 mg/mL of Rifampicin to prevent re-initiation. Reactions were then incubated at 37 °C for 30 sec before seventy-five µL of TRIzol was added to stop the reaction. Nucleic acids were extracted through phenol-chloroform extraction by adding 20 µL of chloroform. Fifty µL of isopropanol, five µL of 5 M sodium chloride, and one µL GlycoBlue (Fisher Scientific, #AM9515) was added to seventy µL of the extracted aqueous phase. The samples were then rehydrated in 43 µL of RNase-free H_2_O. Five µL of DNase TURBO 10X Buffer and 2 µL of DNase TURBO (Invitrogen, #AM2238) were then added, and the reactions incubated at 37 °C for 60 minutes. Reactions were then subjected to an additional round of phenol-chloroform extraction by adding 150 µL of TRIzol and 40 µL of chloroform. The isopropanol precipitation was repeated by doubling the amount of isopropanol and sodium chloride. Pellets were dissolved in 2X loading buffer (8 M urea with bromophenol blue dye (Sigma-Aldrich, #B0126-25G)) and run on a 10% urea PAGE gel at 16 W for 45 min. Gels were stained with Sybr Gold for 5 min and imaged on a BIO RAD ChemiDoc Imaging System. Based on the ssRNA ladder, only the expected terminated and anti-terminated (70 nts and 185 nts, respectively, unless otherwise indicated) band intensities were quantified using Image Lab software from BIO RAD. Percent anti-termination (% AT) was calculated by dividing the intensity of the anti-terminated band by the sum of the terminated and anti-terminated bands. The results were graphed using Datagraph (v4.5.1).

### *Thermus aquaticus* RNA Polymerase and σ^A^ protein purification

The pET28-TaqrpoABCZ^28^ plasmid coexpressing *Thermus aquaticus* (*T. aq*) RNA polymerase four core subunits (α,β,β’ and μ) and the pET28-TaqrpoD^29,30^ plasmid expressing the σ^A^ subunit were co-transformed into *E. coli* BL21(DE3) competent cells. After induction, proteins were purified as previously described^30^.

### DNA template preparation for *T. aq* RNAP *in vitro* transcription

DNA material for *Taq in vitro* transcription first underwent plasmid cloning as previously described to replace introduce the T7 A1 *E. coli* RNA polymerase promoter sequence (ccgaattcaaaaagagtattgacttaaagtctaacctataggatacttacagcc). The double stranded DNA templates were then synthesized through a 200 µL PCR reaction containing 10x Taq Buffer (NEB, #B9014S), 250 µM dNTP Mix (NEB, #N0447L), 125 nM primer C (Supplementary Table S2), 125 nM primer B (Supplementary Table S2), 1 µL DNA plasmid, and 1 µL *Taq* DNA Polymerase (NEB, #M0273X). Reactions underwent a standard thermal-cycle program with 25 cycles of amplifications with an annealing temperature of 53 °C. DNA was purified through an ethanol precipitation with 20 µL 1M NaCl, 1 µL glycogen (Thermo, #AM9515), and 600 µL EtOH. The precipitated pellet was air dried, dissolved in H_2_O, and run on a 2% agarose gel. The template band was extracted using the QIAquick Gel Extraction Kit (Qiagen, #28706X4) and re-purified through ethanol precipitation. DNA concentrations were measured on a Qubit 2.0 Fluorometer (Life Technologies, #Q33216).

### *T. aq* RNAP *in vitro* transcription

Ten µL IVT reactions were assembled with the following concentrations of reagents: 20 mM Tris-HCl pH 8.0, 0.1 mM EDTA pH 8.0, 1 mM fresh DTT, 50 mM KCl, 5 mM MgCl_2_, 0.25 µL BSA, 35 nM DNA template, 100 nM of *T. aq* RNAP, and 200 nM *T. aq* σ^A^. Either 1 mM of NaCl or NaF was also included depending on noted the experimental conditions. Once assembled on ice, the reactions were incubated at 65 °C for 10 min to form transcription complexes.

Transcription was then initiated with the addition of rNTPs to a final concentration of 500 µM, with 125 µM of rATP, rGTP, rUTP, and rCTP. Reactions were then incubated at 65 °C for 10 mins before TRIzol was added to stop the reaction. Nucleic acids were extracted through phenol-chloroform and isolated with isopropanol precipitation. The samples were then rehydrated in 43 µL of RNase-free H_2_O. 5 µL of DNase TURBO 10X Buffer and 2 µL of DNase TURBO (Invitrogen, #AM2238) were then added and the reactions are incubated at 37 °C for 30 minutes. Reactions were then subjected to an additional round of phenol-chloroform extraction by adding 150 µL of TRIzol and 40 µL of chloroform followed by extraction of the aqueous phase. 100 µL of isopropanol was then added to 140 µL of extracted aqueous phase. Pellets were dissolved in 2X loading buffer (8M urea with bromophenol blue dye (Sigma-Aldrich, #B0126-25G)) and run on a 10% urea PAGE gel at 16 W for 45 min. Gels were stained with Sybr Gold for 10 min and imaged on a BIO RAD ChemiDoc Imaging System. Based on the ssRNA ladder, only the expected terminated and anti-terminated band intensities were quantified using Image Lab software from BIORAD by selecting the terminated band. Percent anti-termination (% AT) was calculated by dividing the intensity of the anti-terminated band by the sum of the terminated and anti-terminated bands. The results were graphed using the Datagraph software (v4.5.1).

### Cellular Gene Expression Assays

Plasmids expressing the desired construct (Supplementary Data File S1) were transformed into *E. coli* BW23115 *ΔcrcB*^31^ acquired from the KEIO collection, a strain with the native fluoride transporter knocked out. Three experimental replicates were carried out starting at the transformation step. *E. coli* BW23115 *ΔcrcB* cells were thawed on ice, followed by the addition of 0.2 µL of DNA plasmid and heat shock at 42 °C for 30 second, then incubated at 37 °C at 1,000 rpm in an Incubating Microplate Shaker (VWR, # 97043-606) for 30 minutes. Transformations were grown overnight in a 37 °C incubator on LB agar plates with 34 µg/mL chloramphenicol. The next day the plate was removed from the incubator in the morning and left at room temperature. In the afternoon, three colonies (each a biological replicate) for each construct were selected and deposited in 300 µL LB media with 34 µg/mL chloramphenicol in a 2-mL 96-well block. The cultures were sealed with a Breathe Easier™ Sealing Film (Electron Microscopy Sciences, #70536-20) then shaken overnight at 37 °C at 1,000 rpm in the VWR Incubating Microplate Shaker. The next day, subcultures were made by inoculating 3 µL of overnight culture into 147 µL of freshly made M9 media (1X M9 salts, 1 mM thiamine hydrochloride, 0.4% glycerol, 0.2% casamino acids, 2 mM MgSO_4_, 0.1 mM CaCl_2_) in a new 2-mL 96-well block. The media contained either 500 µM NaCl or 500 µM NaF, unless otherwise stated. These subcultures were sealed with a Breathe Easier™ Sealing Film (Electron Microscopy Sciences, #70536-20) and shaken for 4 hrs at 37 °C at 1,000 rpm unless otherwise denoted.

After the subculture incubation, two parallel measurements were taken (Figure S1). To collect information at a bulk population level, 50 µL of subculture was added to 50 µL of 1X phosphate buffered saline (PBS) with 10 µg/mL of kanamycin in a 96-well optical bottom black plate (Thermo Scientific, #265301). Bulk optical density (OD_600_) and fluorescent readings (excitation = 485 nm, emission = 520 nm, gain = 70) were taken on a Synergy H1 Hybrid Multi-Mode Reader (BioTek, #11-120-535) using Gen5 Microplate Reader and Imager software (v2.04). The arbitrary fluorescent units were normalized by dividing by the OD_600_ before averages and standard deviations were calculated.

Flow cytometry was performed to quantify fluorescence at a single-cell level. Two µL of subculture was added to 200 µL of 1X PBS with 10 µg/mL of kanamycin in a 96-well clear v-bottom plate (Corning, #3896). The samples were run through a BD Accuri^TM^ C6 Plus with a CSampler Plus with the BD CSampler Plus software (v1.0.23.1). Cell events were first gated with an FSC-A threshold set at 2,000. GFP fluorescence was collected on the FL1-A channel monitoring the FITC signal. Data was analyzed using FlowJo software (v10.8.0). An ellipsoid gate on the FSC-A vs SSC-A profile was created to capture *E. coli* cells transformed with an empty plasmid and applied to all datasets. The geometric mean of gated FL1-A fluorescence distribution, to remove outlier peaks as needed, was then extracted (Figure S2A). Fluorescence values were converted to molecules of equivalent fluorescein (MEFL) by calibrating to Sphero^TM^ 8-peak rainbow calibration particles (BD Biosciences, #559123, Lot #8137522). A calibration curve that relates arbitrary fluorescence intensities to MEFL was created by calculating the linear regression between measured relative fluorescence units of each bead distribution and the manufactured supplied MEFL of each distribution. The experimental raw fluorescence was multiplied by the slope of the linear regression curve and the y-intercept was forced to 0 (Figure S2B). The results were graphed using Datagraph (v4.5.1).

### Template-Based Homology Modeling for Molecular Dynamics Simulations

The crystal structure of the holo state *T. pe* fluoride riboswitch aptamer (PDB ID: 4ENC)^8^ was used as a template to model the apo and holo states of the *T. pe* fluoride riboswitch aptamer, as well as the apo and holo states of the *B. ce* fluoride riboswitch aptamer. All modeling was performed using the Molecular Operating Environment (MOE) software package. In MOE, we aligned our *T.pe* and *B.ce* sequences with the crystal structure of the *T.pe* aptamer (PDB ID 4ENC) such that conserved regions were identical and the overall structure was optimally retained. To model the *B. ce* aptamer, the P3 helix of the *T. pe* aptamer crystal structure was reduced by two base pairs in accordance with the predicted structure (Figure 1B). We then mutated the remaining *T. pe* aptamer crystal structure in-place to match the *B. ce* sequence. To model the expression platform, secondary structure diagrams and the sequences of full-length *B. ce* and *T. pe* fluoride riboswitches were used as input into the open-source tertiary structure generator RNAComposer^32^. For both *B. ce* and *T. pe*, RNAComposer predicted 10 possible expression platform conformations. These conformations were narrowed down based on the following criteria: (i) whether the pseudoknot was approximately equidistant from the 3’-end of the expression platform, which competes with the pseudoknot during structural switching, (ii) whether the conformation was in good agreement with the secondary structure diagrams and (iii) whether the backbone path aligned well with that of the existing aptamer models. The chosen expression platform models were graphed onto each aptamer model in MOE. To prevent steric clashes and unrealistic backbone geometry, individual residues near the site where the expression platform was graphed onto the aptamer were energy minimized in MOE. Aside from modeling the apo and holo states of the wildtype *B. ce* and *T. pe* fluoride riboswitches, we additionally modeled the *B. ce* S1 mutant and the *T. pe* W1 mutant by mutating residues of the relevant starting models in MOE.

### All-atom molecular dynamics simulations general overview

All-atom molecular dynamics (MD) simulations were performed using the GROMACS 2019.4 software package^33^. RNA was represented by the Amber99 force field^34^ with Chen-Garcia modifications for RNA bases^35^ and Steinbrecher-Case modifications for backbone phosphates^36^. Mobile magnesium and chloride ions were represented using the Grotz-Schwierz nano-magnesium parameter set optimized for TIP4P-Ew^37^; site-bound magnesium ions were represented using the micro-magnesium parameter set. The interactions between fluoride and site-bound magnesium ions were represented using modified Netz parameters^38^ that were scaled as discussed below (Supplementary Table S2). In holo state models, the magnesium-fluoride complex was modeled using coordinates from the *T. pe* crystal structure. Initial Lennard-Jones parameters for the interaction between fluoride and magnesium were gradually scaled until the tetrahedral geometry of the complex was maintained and the complex remained bound to the ligand binding pocket of the *B. ce* WT holo model over a 16.7 ns MD simulation with distance and angle restraints applied (Supplementary Table S2). These flat-bottomed, harmonic distance restraints were applied at strengths of 250 kJ/mol, along with the angle restraints, between each magnesium ion and fluoride. Additional flat-bottomed harmonic distance restraints were applied at the same strengths to each magnesium ion and the phosphate oxygen of the neighboring nucleotide (Supplementary Table S3). The net effect is that the holo model contains a Mg_3_F ligand with pre-formed ion-RNA contacts that cannot dissociate, whereas the apo model contains mobile Mg^2+^ that are completely free to rearrange. While these assumptions necessarily preclude examination of the ligand binding process itself, it enables the simulations to focus on the downstream structural consequences of the ligand-bound state, in particular, on how it alters favorability of strand exchange. Once prepared, each structure was placed in the center of a cubic box such that there was at least 3.0 nm from all RNA atoms to the box edge in each direction. Structures were solvated with TIP4P-Ew water, 150 mM KCl and 10 mM MgCl_2_. The systems were then energy minimized using the steepest descent algorithm for 1000 steps with a 2 fs timestep and a force tolerance of 800 kJ mol^−1^nm^−1^. Once energy minimized, all systems were simulated in the *N,P,T*-ensemble using a leapfrog Verlet integrator with a time step of 2 fs. An isotropic Berendsen barostat was used for pressure coupling^39^ at 1 bar with a coupling coefficient of Τp = 1 ps, and a velocity-rescale Berendsen thermostat was used for temperature coupling^40^ with a time constant of Τt = 0.1 ps. The LINCS algorithm was used to constrain all bonds, and a cutoff of 1.0 nm was employed using the Verlet scheme for Lennard-Jones and Coulomb interactions. All simulations were run at 310K.

For preliminary simulations of apo *B. ce* that lacked diffuse Mg^2+^, the starting structure was based on the NMR solution structure of the apo modified *B. ce* sequence aptamer structure (PDB ID: 5KH8). This structure was extended in MOE using the secondary structure of the native *B. ce* riboswitch sequence detailed above and previously^23^.

### Equilibration of Homology Models

All homology models were equilibrated for 16.7 ns. During equilibration, flat-bottomed, harmonic distance restraints were applied on capping base pairs and known long-range interactions at strengths of 250 kJ/mol. Since past studies have shown that the apo state of the fluoride riboswitch must access an “excited” state in which the linchpin base pair (A40-U48) is broken to allow the expression platform to disrupt the pseudoknot^8,24,26^, we broke the linchpin during the equilibration of the apo state, but distance-restrained the linchpin during the equilibration of the holo state.

### Simulations of Attempted Structural Switching

To simulate expression platform folding in the *B.* ce and *T.* pe apo and holo state models, flat-bottomed, harmonic distance restraints were applied at strengths of 175 kJ/mol between each base in the 3′-end of the expression platform and the base it pairs with in the pseudoknot during structural switching at strengths of 250 kJ/mol (Supplementary Table S4). The apo and holo states were then each simulated for 40 ns across 20 replicas, for a total of 800 ns per model, and a cumulative total of 6.488 µS of simulation time (Supplementary Table S5). The starting conformations were identical, but the starting velocities were randomized to allow each replica to evolve uniquely. Trajectories were analyzed by tallying the frequency of strand displacement events at each position in the pseudoknot and by calculating the time-averaged radial distribution function, g(r), of mobile magnesium ions with respect to the surfaces of nucleotides in each motif.

## RESULTS

### Establishing assays for fluoride riboswitch function

We first sought to develop assay conditions to characterize fluoride riboswitch function, both *in vitro* and in cells, to validate that functional properties of riboswitch sequence variants observed in minimal *in vitro* systems were reproducible in the complex cellular environment. Previous works used *in vitro* transcription (IVT) to investigate fluoride riboswitch function, using 10 mM NaF to induce switching^22–24^. In our assay conditions, we observed that in the absence of fluoride (1 mM NaCl), the *B. ce* fluoride riboswitch anti-terminated to a level of 28%, which was increased to 75% in the presence of 1 mM NaF (2.68-fold change) (Figure 1C). To demonstrate that this increase in transcriptional read through was due to ligand regulation, we constructed a variant, *B. ce* U41A, in which the highly conserved residue adjacent to the linchpin base pair of the fluoride aptamer is mutated^31^. As expected, *B. ce* U41A was ligand-unresponsive, displaying 16% of anti-termination (AT) independent of added NaF (Figure 1C).

We next sought to investigate riboswitch function through quantitative cellular gene expression assays. Previously, a cell-based assay was developed to demonstrate regulation by the fluoride riboswitch using a β-galactosidase reporter system quantified using a plate-based Miller assay^31^. Here, we sought to develop a non-enzymatic, fluorescence-based cell assay to investigate fluoride riboswitch function at the single-cell level using flow cytometry. To do so, we utilized an expression construct we previously developed to study riboswitch function in cell-free gene expression assays^7^. The expression construct consisted of a *E. coli* sigma 70 promoter, the 79 nt *B. ce* Wild Type (WT) fluoride riboswitch sequence (Figure 1B), a synthetic ribosome binding site, the super-folder GFP^27^ (sfGFP) coding sequence, and a transcriptional terminator (Methods, Supplementary Data File 1). This expression construct was cloned into the p15a plasmid backbone with chloramphenicol resistance. Following previous work, plasmid constructs were transformed into *E. coli* strain BW25113 *ΔcrcB* lacking the CrcB fluoride transporter^31^. Transformants were grown overnight in LB treated with chloramphenicol, and then incubated the next day with increasing NaF concentration in minimal media for 6 hrs (Figure S1).

Flow cytometry was used to quantify single-cell fluorescence for the *B. ce* WT riboswitch expression construct and two negative controls - a constitutively expressed sfGFP expression construct and an expression construct containing the *B. ce* U41A variant (Figure S1B). To quantify the NaF-dependent increases in fluorescence for each construct, we calculated the fold change in fluorescence between each NaF condition and the control (no NaF) for each construct (Figure 1D). For the 1 mM NaF condition, we observed a 5.20-fold increase of fluorescence over control for the *B. ce* WT riboswitch, but also 1.81- and 1.44-fold increases for constitutively expressed sfGFP and *B. ce* U41A, respectively (Figure 1D, Figure S1B). For the 500 µM NaF condition, we observed a 3.26-fold increase of fluorescence for the *B. ce* WT riboswitch, a 1.16-fold for constitutive sfGFP and no change for *B. ce* U41A (Figure 1D, Figure S1B). We therefore chose to proceed with the 500 µM NaF condition.

We noticed that OD_600_ of cell cultures was reduced after 6hrs of growth in 500 µM NaF compared to the no NaF condition (Figure S1C). We therefore performed a time-course experiment in order to find a shorter incubation time that would result in a measurable fold change due to NaF induction and found that fold-changes between 500 µM NaF and 500 µM NaCl growth conditions after four hours of growth to be 2.31, 1.06 and 0.88 for the *B. ce* WT, constitutive sfGFP, and *B. ce* U41A constructs, respectively (Figure S1E). We note that while cell growth in 500 µM NaF is impaired, likely due to fluoride toxicity^41^ (Figure S1E), these assay conditions enable the measurement of changes in gene expression upon NaF inductions for the *B. ce* WT. In addition, the *B. ce* U41A control behaves as expected. However, due to the challenges of cellular assays from NaF toxicity, we decided to utilize both IVT and cellular assays in parallel.

### The pseudoknot sequence influences regulation by the fluoride riboswitch

The fluoride riboswitch architecture consists of mutually exclusive aptamer and expression platform domains that overlap at a key sequence tract – in the aptamer structure this sequence forms a part of the six base pair pseudoknot required for fluoride binding, while in the expression platform this sequence forms the base of the intrinsic terminator hairpin^31^ (Figure 1B). Recent efforts to uncover how fluoride binding regulates the switching mechanism between these two states have revealed that the pseudoknot forms during transcription independent of the presence of fluoride^23,24^. As transcription continues, the expression platform forms into a terminator hairpin that disrupts pseudoknot base pairs. However, the binding of fluoride in the aptamer long-range base pair interactions within the aptamer that prevent the forming terminator from disrupting the pseudoknot (Figure 1A)^22,25^. In other riboswitch systems, it has been observed that sequence context can influence expression platform folding pathways that involve aptamer disruption^15,19^. In particular, strengthening base pairs in portions of the aptamer that must be disrupted has been shown to result in a loss of switching ability or a reduction in dynamic range^15^. Based on this observation, we hypothesized that the composition of the six base pair sequence of the pseudoknot could influence terminator folding and overall fluoride riboswitch function^23,24^.

To test this hypothesis, we explored the effect of sequence variations in the pseudoknot/terminator overlap region (Figure 1B) on riboswitch function. Because prior work demonstrated that mutations in the pseudoknot sequence can break riboswitch function^31^, we constrained our investigations to naturally observed aptamer sequence variation within the Rfam database^42^. Conservation analysis of the annotated fluoride riboswitch aptamer sequences within Rfam 14.9 shows a high degree of conservation in the pseudoknot, which we hypothesized would preclude a systematic exploration of all possible pseudoknot sequence contexts since most of them would not satisfy the natural sequence conservation pattern (Figure 2A). We therefore decided to focus on a natural sequence representative that differ the most from the *B. ce* WT system.

**Figure 2.**
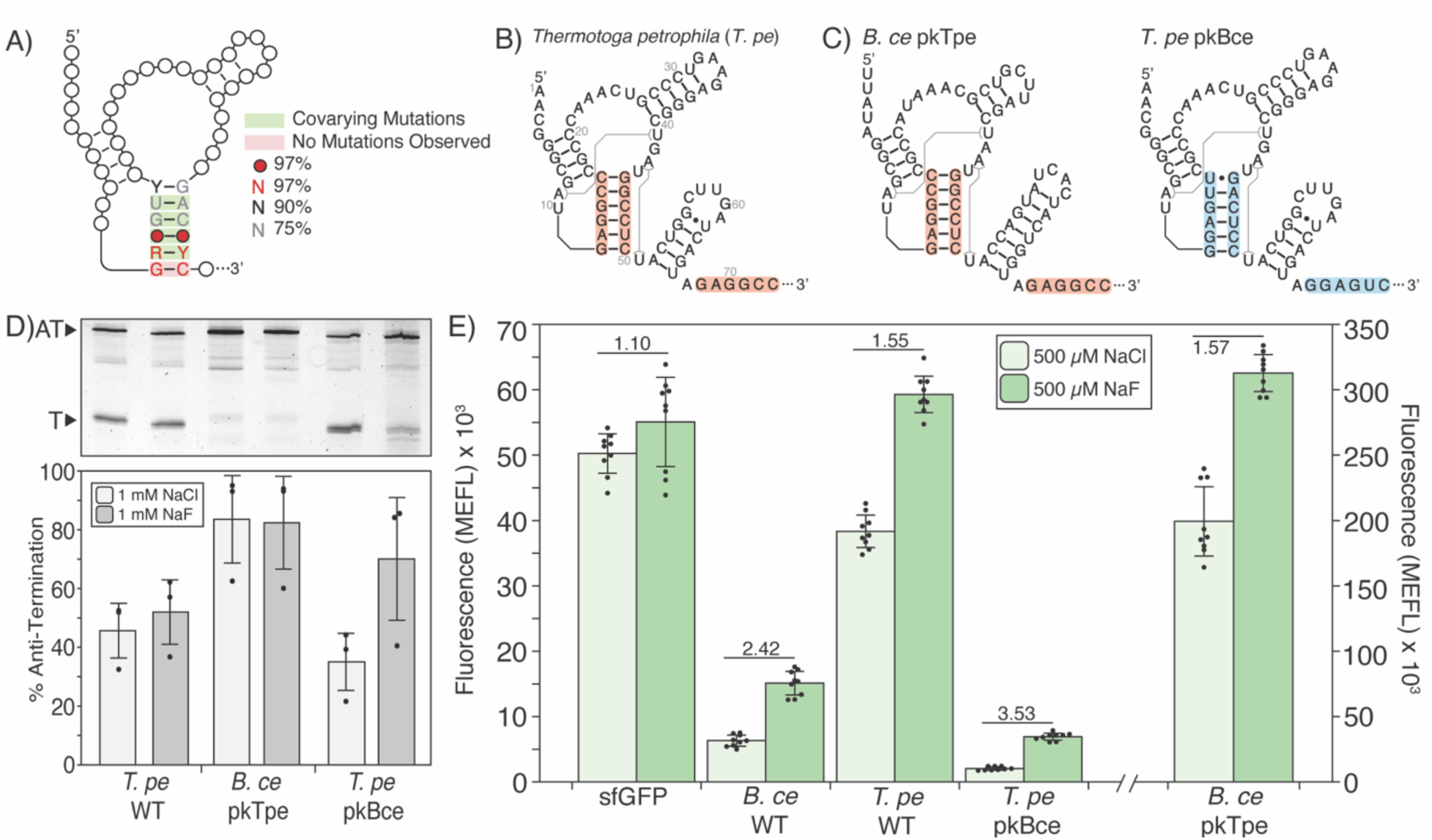
Changes in aptamer pseudoknot sequence alter fluoride riboswitch function. A) 2D representation of the fluoride riboswitch aptamer domain with the evolutionary conservation of the pseudoknot sequence adapted from RFAM (RF01734). B) Holo secondary structure of the *Thermotoga petrophila* (*T. pe*) fluoride riboswitch sequence annotated with aptamer/expression platform overlap sequences. C) Holo secondary structure of chimeric fluoride riboswitches, where the pseudoknot and expression platform overlapping sequence are swapped. *B. ce* with the *T. pe* pseudoknot/overlapping sequences (*B. ce pkTpe*) is on the left, and *T. pe* with the *B. ce* pseudoknot/overlapping sequences (*T. pe pkBce*) is on the right. *B. ce* pseudoknot/overlapping sequences are colored blue with the same *T. pe* sequences colored red. D) Single-round *in vitro* transcription reactions with *E. coli* RNAP for *T. pe* WT, *B. ce* pkTpe, and *T. pe* pkBce riboswitches performed with 1 mM NaCl or 1 mM NaF. A representative gel is shown above a plot of experimental replicates. The Anti-Terminated (AT) (185 nt) and Terminated (T) (70 nt) bands are indicated. E) Cellular gene expression assays of sfGFP control and WT and chimeric *B. ce* and *T. pe* riboswitches subjected to either 500 µM NaCl or 500 µM NaF added to the media. Fold change (NaF condition divided by NaCl condition) is indicated. The y-axis for the *B. ce* pkTpe construct (separated by broken x-axis) is shown on the right. Data in D represents three experimental replicates (N=3), with bars representing the average and error bars representing standard deviation. Replicate gels are found in Supplementary Data File 3. Data in E represent average fluorescence for three biological replicates, each analyzed over three technical replicates (N=9), with error bars representing standard deviation. MEFL – molecules of equivalent fluorescein.

We chose the thermophilic *Thermotoga petrophila* (*T. pe*) sequence, which has an aptamer domain with 83% GC content in the pseudoknot compared to 50% in the *B. ce* pseudoknot, and was used to solve the crystal structure of the fluoride aptamer ^8^ (Figure 2B). We hypothesized that the high GC content of the *T. pe* aptamer would present a barrier to expression platform folding, leading to either a broken switch or reduced dynamic range. To test this hypothesis, we first performed IVT characterization, which showed that the *T. pe* switch antiterminates with ∼50% efficiency, regardless of the presence of fluoride, indicating that the switch is nonfunctional in the assay conditions (Figure 2D).

As a further test, we performed cellular gene expression assays with the *T. pe* WT fluoride riboswitch, which showed some switching, but with a diminished dynamic range of 1.55-fold increase of fluorescence in the presence of fluoride, compared with the 2.42-fold increase of fluorescence by the *B. ce* WT fluoride riboswitch (Figure 2E). Notably, the overall fluorescence of the *T. pe* WT system was higher than that of *B. ce* WT, with the 500 µM NaCl condition resulting in 3.8±0.3 x 10^4^ MEFL compared to 6.3±0.8 x 10^3^ MEFL respectively, and similar to that of the sfGFP control (5.0±0.3 x 10^4^ MEFL) (Figure 2E). One possible explanation for this observation is that the *T. pe* WT terminator is only partially functional *E. coli*. To test this possibility, we created additional control constructs that mutated the aptamer domain to favor terminator formation (Figure S3). Testing two possible constructs with the *B. ce* system revealed that removing the P1 stem (*B. ce*-P1) resulted in similar expression outputs as no fluoride conditions in both IVT and cellular gene expression assays (Figure S3). Characterization of a similar *T. pe*-P1 control in the cellular gene expression assay showed that without the pseudoknot, the *T. pe* terminator is strong and results in a lower fluorescence level than the *T. pe* WT switch without fluoride (Figure S4B), indicating that the terminator is functional. It is possible that the higher overall fluorescence levels observed with the *T. pe* riboswitch in cells is due to the presence of highly stable GC-rich RNA structures within the 5’ untranslated region of the reporter gene mRNA, which are known to stabilize mRNAs and lead to higher levels of expression^43^.

Overall, we found that changing the aptamer pseudoknot sequence context did change functional performance of the fluoride riboswitch.

### Chimeric fluoride riboswitches have behaviors that match the pseudoknot/terminator overlap sequence context

As a further test of the effect of changing pseudoknot sequence on switching function, we created chimeric riboswitches by placing the pseudoknot/terminator overlap region from one riboswitch within the other switch scaffold resulting in two chimeras: *B. ce* pkTpe (*B. ce* with the *T. pe* pseudoknot/terminator), and *T. pe* pkBce (*T. pe* with the *B. ce* pseudoknot/terminator) (Figure 2C). Interestingly in both IVT and cellular conditions, the chimeric riboswitch constructs mimicked the function and phenotype of the riboswitch from which the pseudoknot/terminator overlap sequence originated from. In particular, *B. ce* pkTpe mimicked *T. pe* WT by being non-functional in IVT assay, showing ∼84% AT efficiency independent of fluoride addition (Figure 2D). In cellular conditions, *B. ce* pkTpe had a dynamic range nearly identical to *T. pe* WT (1.57-fold compared to 1.55-fold), and a higher overall elevated level of expression, which, however, was higher than that for *T. pe* WT (Figure 2E). On the other hand, the *T. pe* pkBce chimera was functional in IVT, showing a 2-fold increase in antitermination efficiency, from 35% to 70%, in the presence of fluoride (Figure 2D). The *T. pe* pkBce chimera was also functional in cellular conditions, showing a 3.53-fold increase of fluorescence with fluoride with overall fluorescence levels similar to those of the *B. ce* WT switch (Figure 2E).

Overall, these results show that changing the pseudoknot/terminator overlap sequence has a functional effect on fluoride riboswitch function, supporting our hypothesis that stronger pseudoknot base pairing can lead to diminished dynamic range or broken ON switches.

### Single nucleotide variations within the aptamer pseudoknot alters riboswitch function

We next sought to determine the functional impact of mutating specific positions within the 6 base pairs of the pseudoknot/terminator overlap sequence (Figure 3A, 3B). As discussed above, we constrained this analysis to sequences present within the natural covariation pattern of the pseudoknot (Figure 2A), which reduced the number of possible sequences to investigate.

**Figure 3.**
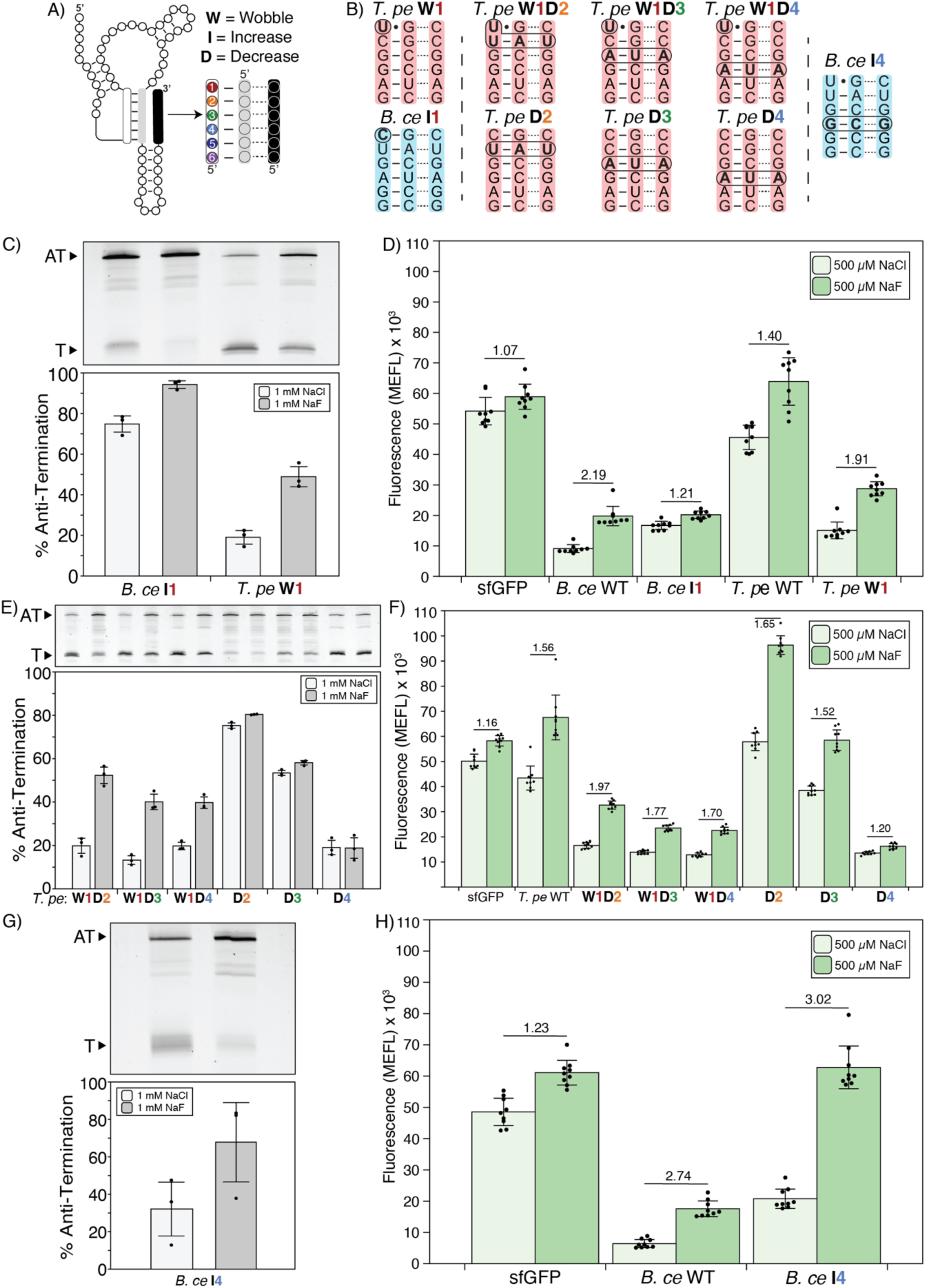
Characterizing the effect of pseudoknot sequence changes on riboswitch function. A) 2D representation of the fluoride riboswitch annotated with mutant naming scheme. Mutation positions use the naming scheme on the right. B) Graphic representation of mutated riboswitches. Mutations are marked through bolding and circling. Mutant sequences are labeled above the mutations and occur within the full sequence context of the indicated *T. pe* or *B. ce* riboswitches. C, E, G) Single-round *in vitro* transcription reactions with *E. coli* RNAP for each mutant fluoride riboswitch performed with 1 mM NaCl or 1 mM NaF. A representative gel is shown under a plot of experimental replicates. The Anti-Terminated (AT) and Terminated (T) bands are indicated. D, F, H) Cellular gene expression assays of the mutant fluoride riboswitches subjected to either 500 µM NaCl or 500 µM NaF added to the media. Fold change (NaF condition divided by NaCl condition) is indicated. Data in C, E, G represent three experimental replicates (N=3), with bars representing the average and error bars representing standard deviation. Replicate gels are found in Supplementary Data File 3. Data in D, F, H represent average fluorescence for three biological replicates, each analyzed over three technical replicates (N=9), with error bars representing standard deviation. MEFL – molecules of equivalent fluorescein.

We focused first on pseudoknot position 1, which is a wobble base pair in the *B. ce* WT fluoride riboswitch but a GC pair in the *T. pe* WT fluoride riboswitch (Figure 1B, 2B). We found that mutating the U to C in the *B. ce* system (*B. ce* I1, where the I denotes an Increase in the number of hydrogen bonds in the base pair) led to a nearly completely anti-terminating riboswitch *in vitro,* with 75% AT in the absence and 94% AT in the presence of fluoride (Figure 3C). Interestingly, testing the inverse mutation (converting the GC pair to a wobble pair) in the *T. pe* system (*T. pe* W1, where the W denotes a wobble base pair), led to a functional switch *in vitro* with 19% AT in the absence and 49% AT in the presence of fluoride (Figure 3C). This swap of function was recapitulated in the cellular gene expression assay, where *B. ce* I1 only had a 1.21-fold increase of fluorescence in the presence of fluoride compared to a 2.19-fold increase observed for *B. ce* WT (Figure 3D). Similarly, *T. pe* W1 showed a greater increase of fluorescence in the presence of fluoride compared to *T. pe* WT (1.91-fold compared to 1.40-fold), with similar overall fluorescence levels to the *B. ce* WT system (Figure 3D). Together these data suggest that the wobble pair at position 1 in the pseudoknot is important for functional switching.

We next sought to investigate how destabilizing base pairs in other positions of the pseudoknot impact riboswitch function. We created a series of *T. pe* fluoride riboswitch mutants where the strength of base pairing between positions 2 and 4 of the pseudoknot was decreased by substituting the GC pairs for an AU pairs. Notably, position 2 in the pseudoknot is evolutionarily conserved to be an UA base pair 75% of the time but is CG in the *T. pe* sequence. Each decreased base pairing strength mutant (D2, D4, where the D denotes the Decrease in the number of hydrogen bonds in the base pair) was tested in the context of the wobble (W1) or the natural GC pair in position 1 (Figure 3B). In IVT, we found that the D2 and D4 variants were non-functional and did not respond to 1 mM NaF, while the mutants that also contained the wobble in position 1 (W1D2, W1D4) were functional switches (Figure 3E). The cellular assay showed similar trends as before, with the *T. pe* W1D2 and W1D4 variants showing both fluorescence levels and switching dynamic ranges similar to *B. ce* WT, while the *T. pe* D2 variant showing elevated overall fluorescence and reduced dynamic range similar to *T. pe* WT.

To further explore the observation that even weakened *T. pe* pseudoknot variants needed the wobble base pair in the first position to switch, we constructed weakened third position variants, since the 5^th^ position is already a weakened base pair. We found *T. pe* D3 variant anti-terminated about 60% of the time regardless of fluoride presence but introducing the wobble base pair resulted in a functional riboswitch (Figure 3E, 3F). This further supports the importance of the wobble base pair.

Interestingly, the mutant *T. pe* D4 anti-terminated about 20% of the time regardless of fluoride presence both within IVT and cellular conditions, which could reflect that this position affects the switching process differently than the other positions investigated. To further investigate this, we introduced a corresponding mutation to the *B. ce* system by increasing the strength of the base pair in position 4 from an AU to a GC pair (Figure 3B). The mutation had no impact the fluoride riboswitch *in vitro* function (Figure 3G) compared to *B. ce* WT (Figure 1C). Additionally, the cellular gene expression assay revealed that *B. ce* I4 had a similar fold-change as *B. ce* WT, however with an increase of baseline fluorescence (Figure 3H).

Taken together, these results demonstrate that the base pairing composition of position 1 of the pseudoknot has the most influence on fluoride riboswitch function, and in particular that a wobble base pair is needed for switching under these assay conditions.

### The pseudoknot wobble pair influences fluoride riboswitch function even at elevated temperatures

The above results show that weakening the *T. pe* fluoride riboswitch in key positions can lead to functional switching at 37 °C, and suggests that the lack of switching for the wild-type system at these temperatures could be due to the high GC content of the pseudoknot, which prevents for structural switching. However, the *T. pe* fluoride riboswitch sequence evolved in a thermophilic environmental condition at elevated temperatures (47 – 88 °C)^44^, raising an important question as to whether increasing the temperature above the conditions already investigated can improve the function of the wild type riboswitch. To investigate this, we adapted our IVT assay to function at elevated temperatures by substituting the *E. coli* RNAP with the *T. aq* RNAP, previously used to investigate the effect of temperature on transcription initiation^29,45–48^, and transcription termination^29^.

Assay conditions with *T. aq* RNAP were based on the *E. coli* single-round IVT assay and modified to increase RNA yield given the weakened RNAP-promoter interactions of *T. aq* RNAP compared to *E. coli* RNAP^45^ by removing the re-initiation blocker rifampicin, conducting transcription for 10 min, and reducing the reaction volume from 25 µL to 10 µL. In addition, the *E. coli* sigma 70 promoter was replaced with the T7A1 promoter sequence. We observed that the *T. pe* fluoride riboswitch terminated less in the absence of fluoride, showing 80% AT at 65 °C (Fig. 4A) compared to 45% AT at 37 °C (Fig. 2D). However, this lack of termination was not due to the lower efficiency of termination by *T. aq* RNAP observed earlier^29^, as the *T. pe*-P1 control showed robust 37% termination (Fig. S6A). For the *T. pe* WT, the observed 80% AT efficiency in the absence of fluoride increased to 94% in the presence of fluoride (Fig. 4A), indicating that indeed this riboswitch is more functionally active at elevated temperatures compared to the complete lack of switching observed at 37 °C (Fig. 2D).

**Figure 4.**
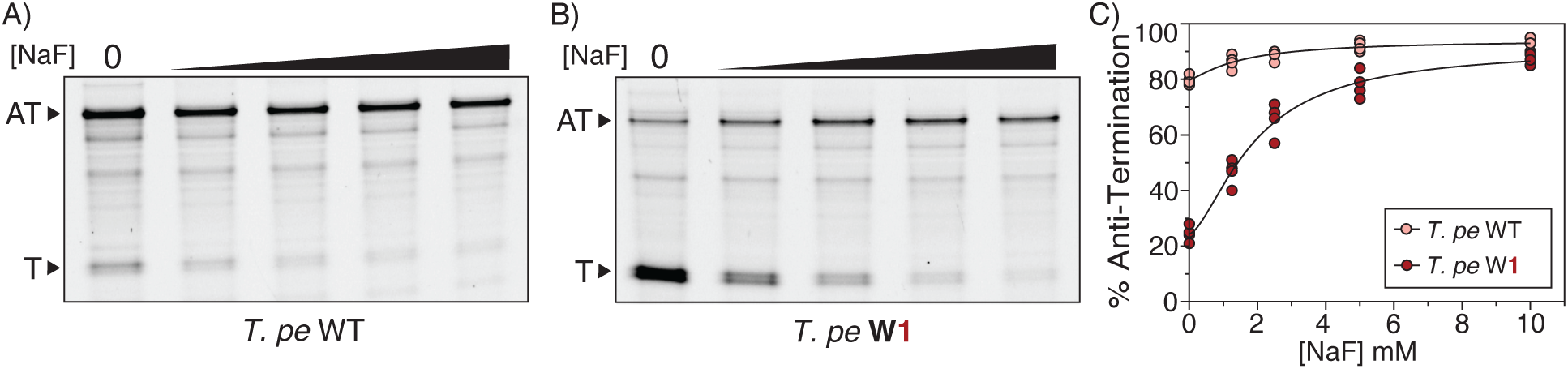
High-temperature transcription of the *Thermotoga petrophila* fluoride riboswitch wild type and wobble base pair variants using *T. aq* RNA polymerase. A) Dose-response *in vitro* transcription reactions with *T. aq* RNAP at 65 °C for *T. pe* WT in the presence of 0, 1.25, 2.5, 5, or 10 mM NaF. A representative gel is shown. The Anti-Terminated (AT) (188 nts) and Terminated (T) (85 nts) bands are indicated. B) Dose-response *in vitro* transcription reactions with *T. aq* RNAP at 65 °C for *T. pe* W1 in the presence of 0, 1.25, 2.5, 5, or 10 mM NaF. A representative gel is shown. The Anti-Terminated (AT) (188 nts) and Terminated (T) (85 nts) bands are indicated. C) A plot of % anti-termination calculated from band intensity quantification of experimental replicates (N=4). Points were fit with nonlinear regression curves (Figure S6B). Replicate gels are found in Supplementary Data File 3.

Given that the *T. pe* riboswitch can function at elevated temperatures, we next sought to test whether the wobble mutation to the pseudoknot would have a similar effect to enhance function at these temperatures. IVT characterization of the *T. pe* W1 variant at 65 °C revealed 24% AT in the absence of fluoride, and 91% in the presence of 10 mM fluoride (Fig. 4B), showing improved dynamic range. Comparison of IVT induction curves of the *T. pe* WT and W1 variants across multiple fluoride conditions shows enhanced dynamic range for the W1 mutation across concentrations, with a similar concentration of NaF to reach half-max anti-termination (AT_50_) for each variant of 1.5 mM for WT and 1.9 mM for W1 (Fig. 4C, S6B), which is an order of magnitude higher than previously seen with *B. ce* WT with the *E. coli* RNAP^22^.

This data expands our findings to show that elevated temperatures do improve the function of the *T. pe* fluoride riboswitch, while the impact of the wobble base pair in the pseudoknot to increase dynamic range is still present at higher temperatures.

### All-atom molecular dynamics simulations reveal how the pseudoknot sequence regulates structural switching

The functional mutagenesis data presented above suggests that structural switching in the fluoride riboswitch is not solely dictated by the linchpin base pair^8,9,11^. In particular, we found that the identity of the wobble base pair at the top of the pseudoknot is a key determinant of riboswitch function. This observation motivated a more detailed investigation of the fluoride riboswitch switching process in order to understand how base identity in the pseudoknot helps regulate switching.

To probe the structural switching process at a more mechanistic scale, we conducted all-atom molecular dynamics (MD) simulations of the apo (ligand unbound), and holo (ligand bound) states of both the *T. pe* and *B. ce* fluoride riboswitches. Since the structure of a full-length fluoride riboswitch has not been solved, we used the crystal structure of the *T. p*e fluoride aptamer^8^ as a template to generate homology models of the apo and holo states of both the *B. ce* and *T. pe* fluoride riboswitches (Figure 5A, 5B). To model the ligand bound holo state, we incorporated three site-bound Mg^2+^ ions within the holo state *B. ce* and *T. pe* models that were distance restrained to both the fluoride ion and three phosphate oxygens within the aptamer, in accordance with the crystal structure (see Materials and Methods). All models were equilibrated for 16.7 ns at 310K. We additionally adjusted the linchpin base pair distance depending on the ligand state: the holo state was equilibrated with the linchpin base pair formed under a distance restraint, while the apo state was equilibrated with the linchpin broken, as previous studies^24,49^ found that the apo state must access an excited conformation where the linchpin base pair is broken to undergo structural switching^24^.

**Figure 5.**
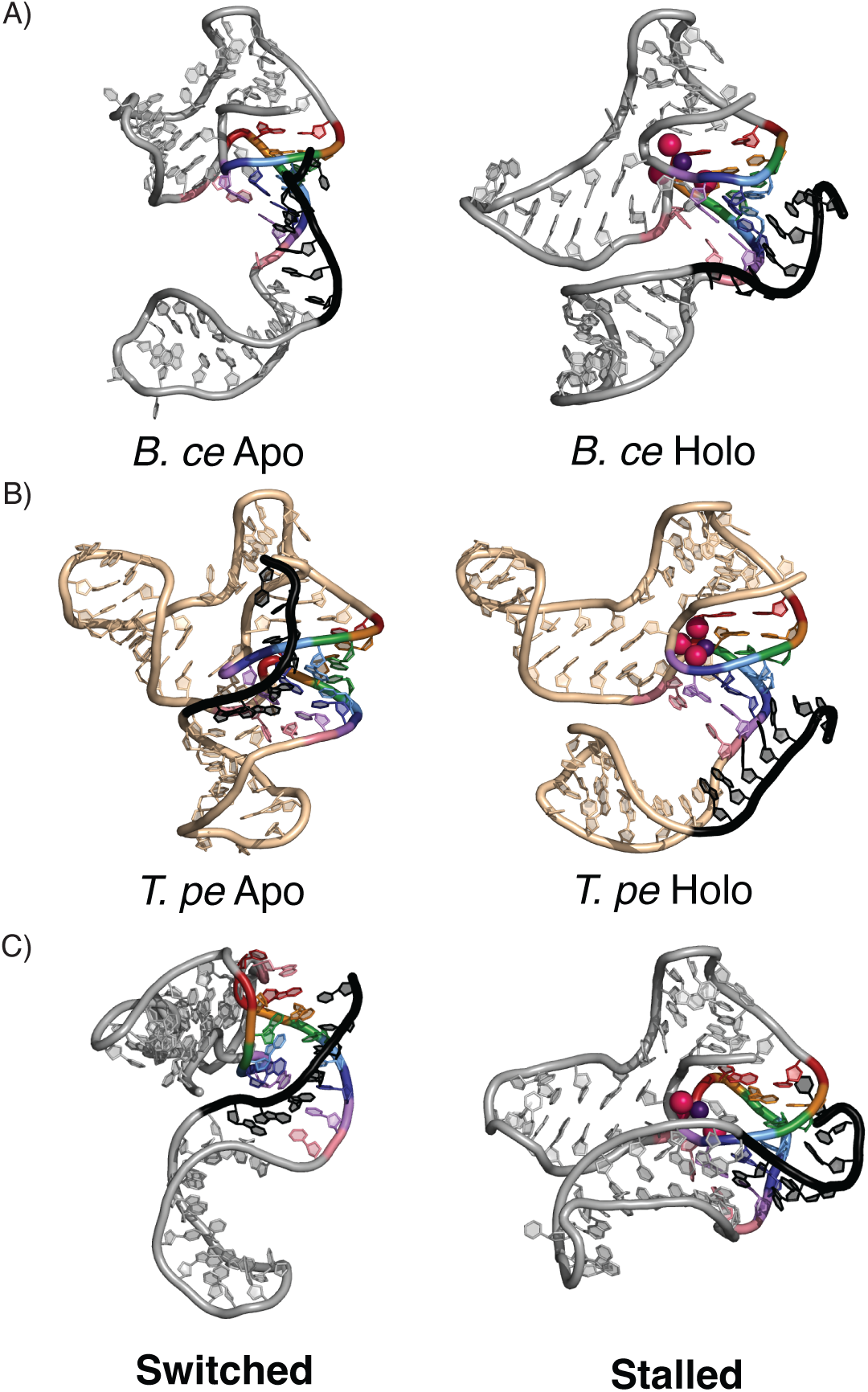
Snapshots of all-atom MD simulations of the switching process in the *B. ce* and *T. pe* systems. Template-based homology modeling of the fluoride riboswitch was based on the *T. pe* aptamer crystal structure to simulate structural switching by the expression platform through the pseudoknot of each riboswitch system. Color coding follows Figure 1A, with the expression platform overlap region colored black. (A) Starting apo and holo states of the *B. ce* system. (B) Starting apo and holo states of the *T. pe* system. (C) An ending conformation of one simulation of the switching process that was successful in the *B. ce* system, where all pseudoknot base pairs were displaced by the expression platform (left), and an ending conformation of one simulation of a stalled switching process in the *B. ce* system, where the bases could not rearrange due to the irreversibility of the biased simulations.

We simulated the structural switching process by applying distance restraints between bases on the invading expression platform strand and their complements on the 3′-end of the pseudoknot (see Materials and Methods). The use of biases to encourage structural switching results in non-equilibrium simulations in which models either undergo successful switching on sub-40 ns timescales, or irreversibly stalling without undergoing switching (Figure 5C)^16^. Since each trajectory samples a single structural switching attempt, each model was simulated for 40 ns across 20 replicas with identical starting conformations but randomized, thus unique, starting velocities. Due to the irreversibility of biased simulations, once a trajectory stalled, it was impossible for bases to rearrange such that switching could be attempted again. For that reason, longer simulations would not provide useful information. Since the same biases were applied to both the apo and holo states, differences in structural switching between the apo and holo states indicate an influence of fluoride-mediated magnesium:RNA interactions on the switching process.

Each trajectory was analyzed for the completion of switching at each base pair in the pseudoknot, measured as the number of hydrogen bonds formed between each substrate and invading base, across 20 replicas for each sequence in the apo and holo states (Figure 5). As expected, in the holo state *B. ce* WT model, switching is rarely seen at each base pair, with only one successful structural switching event that displaced all pseudoknot positions. In contrast, in the apo state, displacement was more frequent, with each base pair displaced at least 35% of the time (Figure 6A). These data are in accordance with the functional data, where the *B. ce* WT fluoride riboswitch in the presence of fluoride still showed some termination in IVT conditions (Figure 1C), where termination is synonymous with complete structural switching.

**Figure 6.**
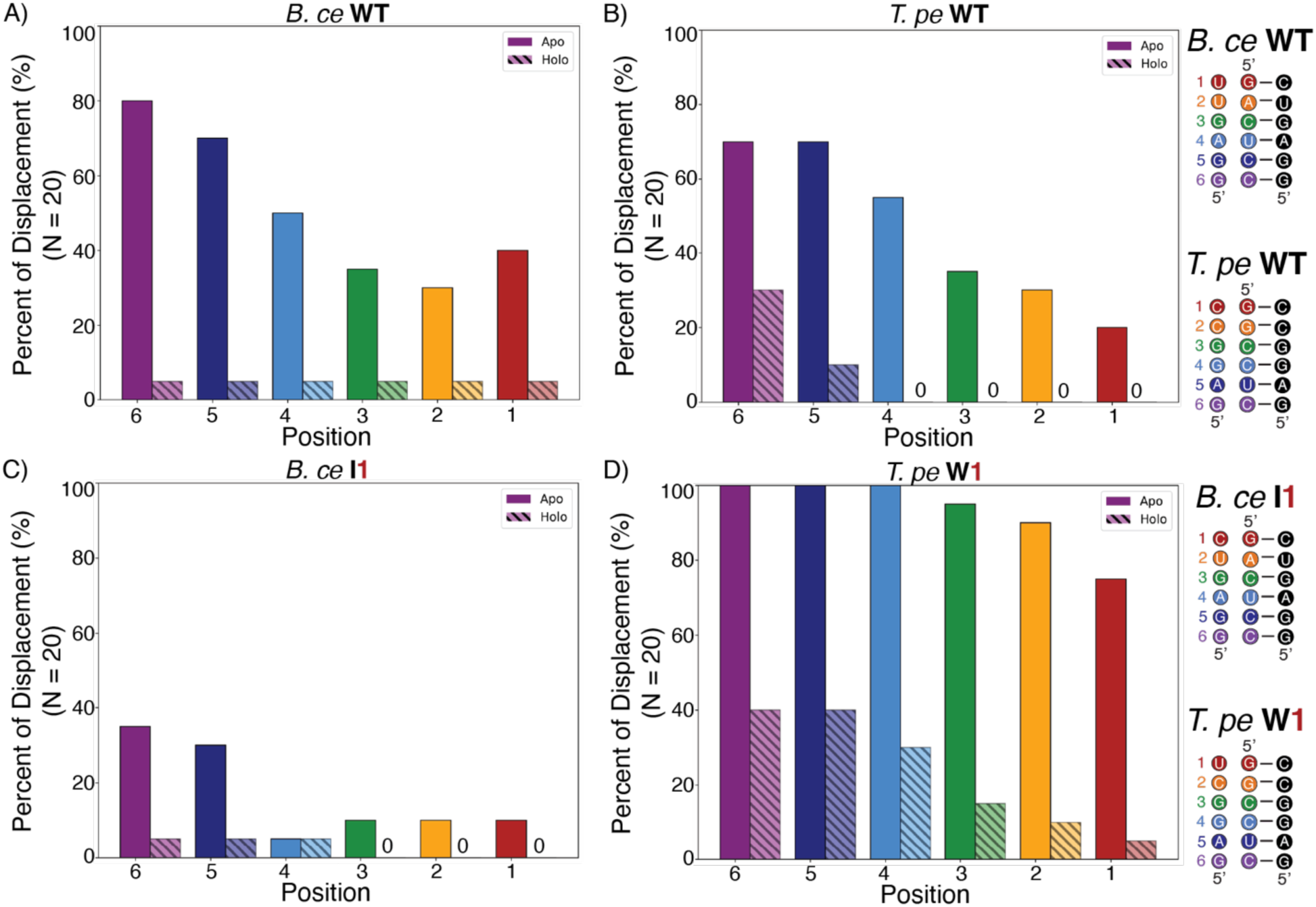
Simulated structural switching of each base pair in the fluoride aptamer pseudoknot by the expression platform. Each of the twenty replica structural switching simulations (N=20) of the apo and holo states for the (A) *B. ce* WT, (B) *T. pe* WT, (C) *B. ce* I1 variant, (D) *T. pe* W1 variant fluoride riboswitch were analyzed to quantify the percentage that each pseudoknot position was displaced by expression platform folding at the end of the simulation. Bars are colored according to their position along the pseudoknot (Figure 1A). Apo state simulations are represented by solid bars and holo state simulations are represented by bars with diagonal lines. The reference sequence for each variant, showing the pseudoknot base pairs and the overlapping expression platform sequence, is provided on the right.

In contrast, the *T. pe* WT model in the holo state never experiences complete structural switching in simulations. In the *T. pe* WT apo state, successful structural switching through the entire pseudoknot only occurred 15% of the time, though some simulations showed partial switching through the positions closer to the expression platform sequence (Figure 6B). This is different than what is observed in IVT conditions, where the *T. pe* WT fluoride riboswitch anti-terminated ∼50% of the time independent of fluoride conditions, showing that some structural switching can occur in this system, though it is not fluoride dependent (Figure 2D). This discrepancy could be attributed to unknown interactions between the riboswitch and RNA polymerase that are only beginning to be understood^50^ and therefore cannot be captured by MD simulations.

Next, we modeled the switching process of the apo and holo states of the *B. ce* I1 and *T. pe* W1. In accordance with the functional data, the simulations of the *B. ce* I1 model show overall less strand displacement compared to *B. ce* WT for every nucleotide in the apo state and no successful complete strand displacement events in the holo state (Figure 6C). In contrast, our simulations of the *T. pe* W1 model show increased strand displacement at each position, with the 6^th^, 5^th^, and 4^th^ positions reaching 100% strand displacement across all simulations of the apo state (Figure 6D). There are also successful strand displacement events in the *T. pe* W1 holo state (Figure 6D), which was observed experimentally as in the presence of fluoride the amount of anti-termination and fluorescence was decreased from the *T. pe* WT system (Figure 3C, 3D).

Apart from comparing the structural switching simulations with the functional results, we analyzed the trajectories to understand how structural switching can occur in a three-dimensional (3D) space. Unlike what appears on a secondary structure diagram, the default conformation of the 3′-end of the expression platform is twisted ∼ 90° from the 3′-strand of the pseudoknot which it must pair to for a complete invasion (Supplementary Figure S7, Supplementary Movies). Importantly, the architecture of the P1 stem-loops prevent the invading strand from achieving a pitch similar to the pseudoknot 3′ strand without severe steric overlap with the rest of the RNA. For this reason, in many trajectories where stalling has occurred, the riboswitch adopts a pseudo-triple helix due to distance restraint biases, and the expression platform strand cannot successfully pair with the pseudoknot 3′ strand. Even in many trajectories where structural switching is successful, the pseudoknot 3′ strand does not completely dissociate from this triple helix structure due to the amount of twisting. In fact, a “clean” structural switching event only occurs if base pairs in the pseudoknot break from the top down, enabling the pseudoknot 3′ strand to rotate and match the pitch of the 3′-end of the expression platform. This explains why out-of-order structural switching is occasionally seen in our trajectories - formation of the top base pair between the expression platform and pseudoknot 3′ strands enables the expression platform strand to reorient itself, thereby “rescuing” the structural switching process that otherwise would have stalled. This could provide a mechanistic explanation as to why the top position in the pseudoknot is consequential for regulating structural switching – if the top position is a wobble base pair (*B. ce* WT or *T. pe* W1), base pairs in the pseudoknot will more easily break from the top down, relieving stress on the structural switching interface by permitting the rotation of the pseudoknot’s 3′-strand.

### All-atom molecular dynamics simulations reveal the role of magnesium colocalization in regulating structural switching

Central to the fluoride riboswitch aptamer holo state are three magnesium ions that coordinate the fluoride ion within the negatively charged RNA structure^8^. To study potential roles of these magnesium ions in the switching process, we utilized recently developed novel magnesium parameters for RNA folding in our simulations^37^ to explicitly model both the coordinated fluoride-Mg^2+^ complex, as well as free Mg^2+^ ions in solution that may associate with the RNA. Interestingly, even with strong bias forces, we were unable to observe complete strand exchange events in simulations that lacked diffuse Mg^2+^ ions in apo riboswitch simulations (Supplementary Figure S8). This strongly suggests that Mg^2+^ plays a more nuanced role in the structural switching process than merely chelating the F^-^ ligand as observed in the crystal structure. To quantify this, we calculated the radial distribution function, g(r), of mobile magnesium ions, with respect to the surfaces of nucleotides in each structural motif of the *B. ce* WT fluoride riboswitch (Figure 7) (see Materials and Methods). In holo state trajectories, due to the presence of site-bound magnesium ions in the ligand binding pocket, more mobile magnesium ions colocalize to the bottom of the pseudoknot, the 3’-end of the expression platform, the P1 helix, and the J1/2 hinge, providing additional stability to these motifs and thereby decreasing the likelihood of structural switching. In contrast, the magnesium radial distribution functions are different in successful apo state trajectories that switched, showing a more delocalized profile of magnesium location that reflects magnesium ions that remain highly diffusive throughout the entire simulation. Interestingly, in failed apo state trajectories that stall without completing structural switching, the same trend as the holo trajectories is observed, with g(r) profiles that show an increased density near the analyzed motifs. This is due to the stochastic accumulation of mobile magnesium ions in the ligand binding pocket, which leads to the colocalization of other mobile magnesium ions to the same motifs, causing the apo state to more closely resemble the holo state.

**Figure 7.**
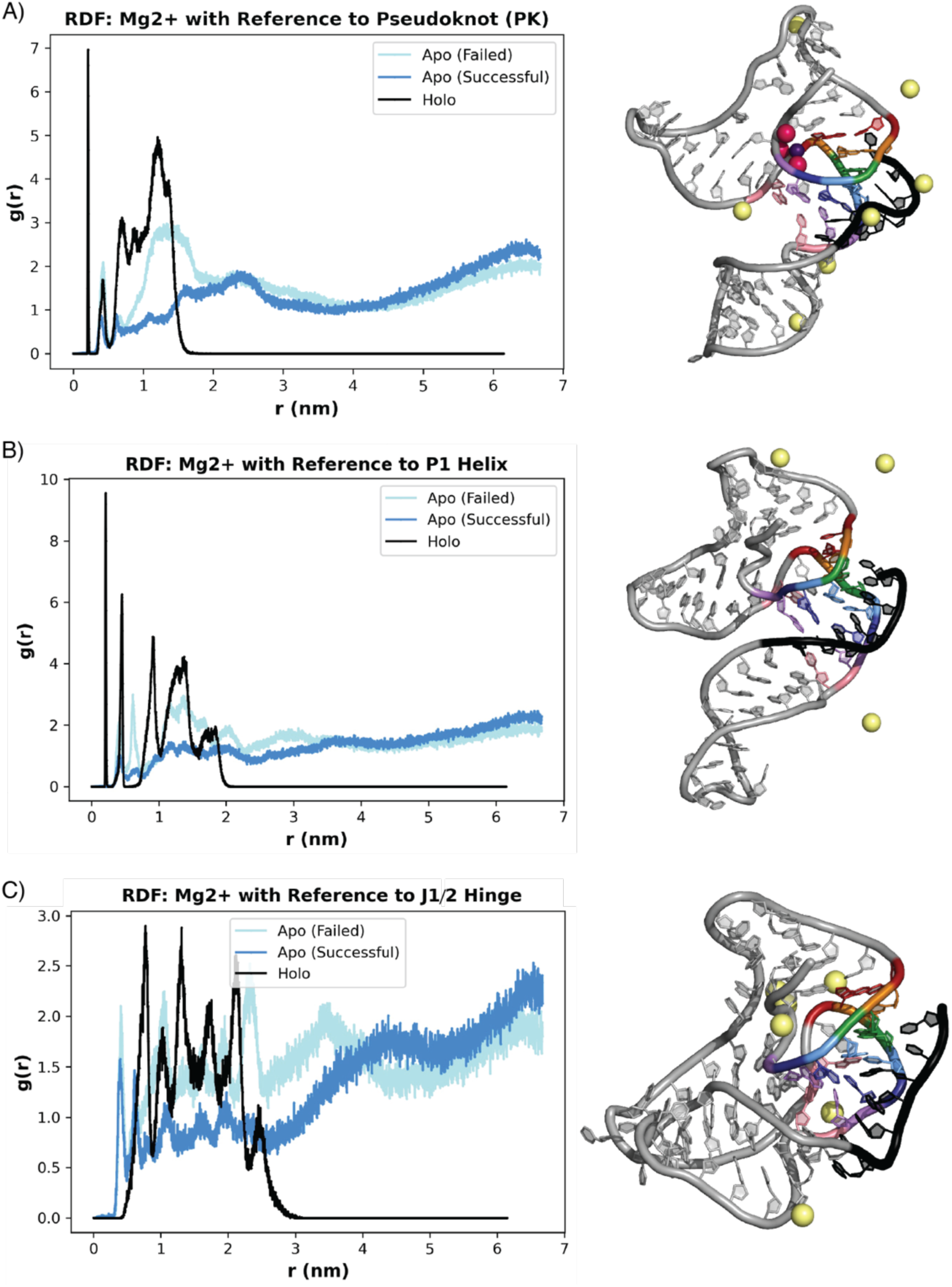
Radial distribution of bulk magnesium around the binding pocket and first position of the *B. ce* pseudoknot. Snapshots from simulation trajectories showing how mobile magnesium ions colocalized to the *B. ce* WT fluoride riboswitch in the holo state (black), the apo state for a replica that underwent successful structural switching (dark blue), and the apo state for a replica that could not undergo structural switching (light blue). Magnesium colocalization patterns are shown as radial distribution functions (RDFs) of magnesium density calculated with respect to (A) the pseudoknot, (B) the P1 helix and (C) the J1/2 hinge. RDFs are calculated from representative traces from single simulation trajectories.

Overall, these simulation results suggest that magnesium ion distribution, specifically colocalization near the aptamer motifs that participate in the switching process, could influence whether successful switching occurs, contributing an additional source of stochasticity to the switching process.

## DISCUSSION

In this work we investigated the influence of aptamer/expression platform sequence on the switching process of the fluoride riboswitch. Previous work investigating the folding mechanism of the fluoride riboswitch primarily focused on the formation of the long-range A40-U48 linchpin base pair as the dominant determinant of whether switching can occur in the fluoride riboswitch^24,26,49^. Our results show that additional factors can also influence switching, in particular the sequence of the aptamer pseudoknot that overlaps with the base of the expression platform terminator (Figure 2). Specifically, our results show the importance of having a wobble base pair at the top of the pseudoknot to allow functional switching of the riboswitch in both IVT and cellular conditions (Figure 3). Our results also show switching remains influenced by the base pair composition even at elevated temperature (Figure 4).

Our functional data was further supported by all-atom MD simulations (Figure 5) that suggested a mechanistic explanation of how the presence of this wobble pair could allow the relief of a topological twist in the switching process (Figure S7), allowing the pseudoknot 3’ end to orient itself for productive pairing to the expression platform region through breaking of the pseudoknot from the top down. This result suggests the importance of considering the three-dimensional geometry of the aptamer/expression platform architecture in understanding the fluoride riboswitch switching process. In particular, considering the three-dimensional geometry helps understand the potential for base pair identity to influence the switching process through three dimensional geometrical changes and not just through free energy differences that are the typical focus of analysis when considering only secondary structures. We further find that, even when strong biases encouraging strand exchange were implemented, strand exchange was only observed in presence of diffuse Mg^2+^ ions (Figure 7, S8), implicating a more nuanced role played by Mg^2+^ beyond simple stabilization of the fluoride at the binding site, in that is also facilitates the necessary three-dimensional contortion of the invading strand required for successful strand exchange.

Our findings demonstrating the importance of the wobble base pair at the top of the pseudoknot of the fluoride riboswitch complements previous observations of an NMR hydrogen exchange experiment^26^. This NMR study revealed that the same aptamer wobble base pair of the *B. ce* WT fluoride riboswitch modified to stabilize the structure for NMR analysis had almost twice the base-pair fluctuations when F^-^ was not present. Specifically, they observed that regardless of F^-^ presence, U17 is less stable and more dynamic even while base paired, compared to its partner G42. They also observed that the wobble base pair openings reduce two-fold when F^-^ is present. This provides additional data to help explain the importance of the aptamer wobble pair in the fluoride switching mechanism, pointing to the role of base pair fluctuation dynamics in this region that can facilitate switching, potentially through the relief of three-dimensional twist as found in our MD simulations.

The tight structural dynamics inhibiting switching appeared to still be present at the elevated temperature *T. pe* is naturally found. To our knowledge, this is the first functional IVT characterization of a riboswitch at a thermophilic temperature. Previously, a translational fluoride riboswitch was demonstrated as a fluoride-responsive inducible system in a cellular assay with archaeal hyperthermophiles^51^. An smFRET study of the manganese riboswitch between 22.2 – 28 °C found that increased temperature could increase RNA folding^52^, which might explain why the *T. pe* WT fluoride riboswitch anti-terminated *more* at the elevated temperature. There may also be additional nucleotide interactions occurring to explain the increased structure stability as seen with NMR studies of the preQ_1_ riboswitch form *Thermoanaerobacter tengcongensis* at 50 and 65 °C^53^. The preQ_1_ riboswitch was also studied through smFRET study, which similarly to our study compared meso- and thermophilic sequences for a preQ_1_ riboswitch and found that the thermophilic variant is more thermostable and that solvents impacted each sequence differently^54^. Further studies comparing RNA sequences with similar functions but evolved in different environments may elucidate other adaptive genomic sequence features. Our thermophilic IVT assay adds to the growing body of techniques to investigate the role of temperature on RNA structure dynamics.

All-atom MD also allowed us to also investigate beyond the structural features to the role of magnesium ions in the structural switching mechanism. While the role of magnesium in helping to establish stable RNA folds has long been appreciated^55,56^, much less is known about the role of magnesium in facilitating RNA folding dynamics and rearrangements. Previous all-atom MD studies showed the ability of magnesium to influence RNA structure around the ligand binding pocket of the *add* adenine riboswitch^57^. In this work, our all-atom MD simulations of the folding of the fluoride riboswitch suggest an interesting dynamic between the fluoride ligand and magnesium folding cofactor. In particular, we observed that the site-bound Mg_3_F complex indirectly influences both local RNA flexibility in the ligand-binding region and mobile Mg^2+^ occupancies through Mg^2+^ colocalization patterns (Figure 7). Specifically, radial distribution functions show that mobile Mg^2+^ ions are concentrated at the structural switching interface in the holo state when the Mg_3_F complex is present (Figure 7). This additional concentration of mobile magnesium ions appears to help prevent switching, as we observed a similar distribution of ions in apo state trajectories that did not successfully switch (Figure 7). Intriguingly, we were unable to observe any structural switching in extensive preliminary MD simulations (Supplementary Figure S8) of the *B. ce* WT fluoride riboswitches in the apo state using similar biasing forces but only monovalent ions. Given the apparent influence of Mg^2+^ on the switching mechanism of the fluoride riboswitch, it could be argued that the riboswitch senses both fluoride and magnesium, though the independent effect of magnesium on switching would be difficult due to the influence of magnesium concentration on transcription^58^.

Previous work on the fluoride riboswitch mechanism suggested a role for the cellular transcription factor NusA in the cotranscriptional folding of the riboswitch^22^. These IVT studies found that the cellular factor NusA increased termination rate in the absence of fluoride and smFRET results show that NusA continuously binds and unbinds the nascent transcript. In the presence of fluoride, the stabilized RNA structure results in NusA release. In our work, we observe that some sequences (*T. pe* WT, *B. ce* pkTpe, *T. pe* D2, *T. pe* D3) show slight to no fold-change in the *in vitro* assay have some functionality in cells. As NusA was shown to increase termination in the absence of the fluoride riboswitch, it could be that the slight functionality we observe in cells is the impact of NusA, or other cellular factors, rather than fluoride binding decreasing termination.

This work fits into several studies that have revealed the influence of small sequence changes on cotranscriptional RNA structural rearrangements^15,16,19^. In other riboswitch systems such as the ZTP riboswitch, single point mutations to the aptamer/expression platform overlap sequence were shown to tune the riboswitch dynamic range through influencing the kinetics of a strand displacement process^17^. In the *yxjA* adenine riboswitch, single base pair changes were shown to completely bias the folding pathway towards the riboswitch ON or OFF state again through influencing the competition between a strand displacement switching mechanism^16^. In the translational Guanidine-II riboswitch, researches quantitatively explored every single point mutant impact on riboswitch function^18^. They similarly found that point mutations could impact riboswitch dynamic range, sensitivity, and that specific mutations in the non-conserved linker region could break riboswitch function likely by changing the conformational landscape. In the Signal Recognition Particle (SRP) RNA, a wobble base pair has been shown to be critical for allowing a cotranscriptional structural switching process to occur through allowing hairpin base pair fluctuations needed for strand invasion^59^. Here we add another potential way through which single base pair changes can influence RNA structural switching, by having a wobble pair mediate RNA fluctuations that allow proper three-dimensional alignment of strands that must pair with each other to switch.

Overall, this work adds to the growing understanding of the sequence determinants that facilitate dynamic RNA structural switching at the heart of riboswitch mechanisms. Our all-atom MD simulations also provide additional insight into how magnesium can influence these RNA folding dynamics. While our study has been focused on the fluoride riboswitch, we anticipate these findings to be useful for understanding how RNA base pair changes could affect RNA-mediated diseases through affecting RNA structural ensembles^60–63^, and also how to program RNA dynamics within RNA biotechnologies^64^.

## AUTHOR CONTRIBUTIONS

Conceptualization: L.C., A.M.Y, L.M.H., A.C., J.B.L. *E. coli in vitro* and cell experiments: L.M.H. *T. aq in vitro* assays: L.M.H., K.K. Simulations: E.N.W., R.K. Funding acquisition: A.M.Y., L.M.H., A.C., J.B.L. Writing (original draft): L.M.H. Editing: all authors.

## ACKNOWLEDGMENTS

We thank Katherine Berman for training L.M.H. on experimental techniques and Walter Thavarajah for helping to develop the *T. aq* IVT assay. L.M.H. thanks Yvette Marie Babineau (1962-2020) for personal and professional support.

