## Supplementary Information for "The Effect of Pseudoknot Base Pairing on Cotranscriptional Structural Switching of the Fluoride Riboswitch"

### Figure S1

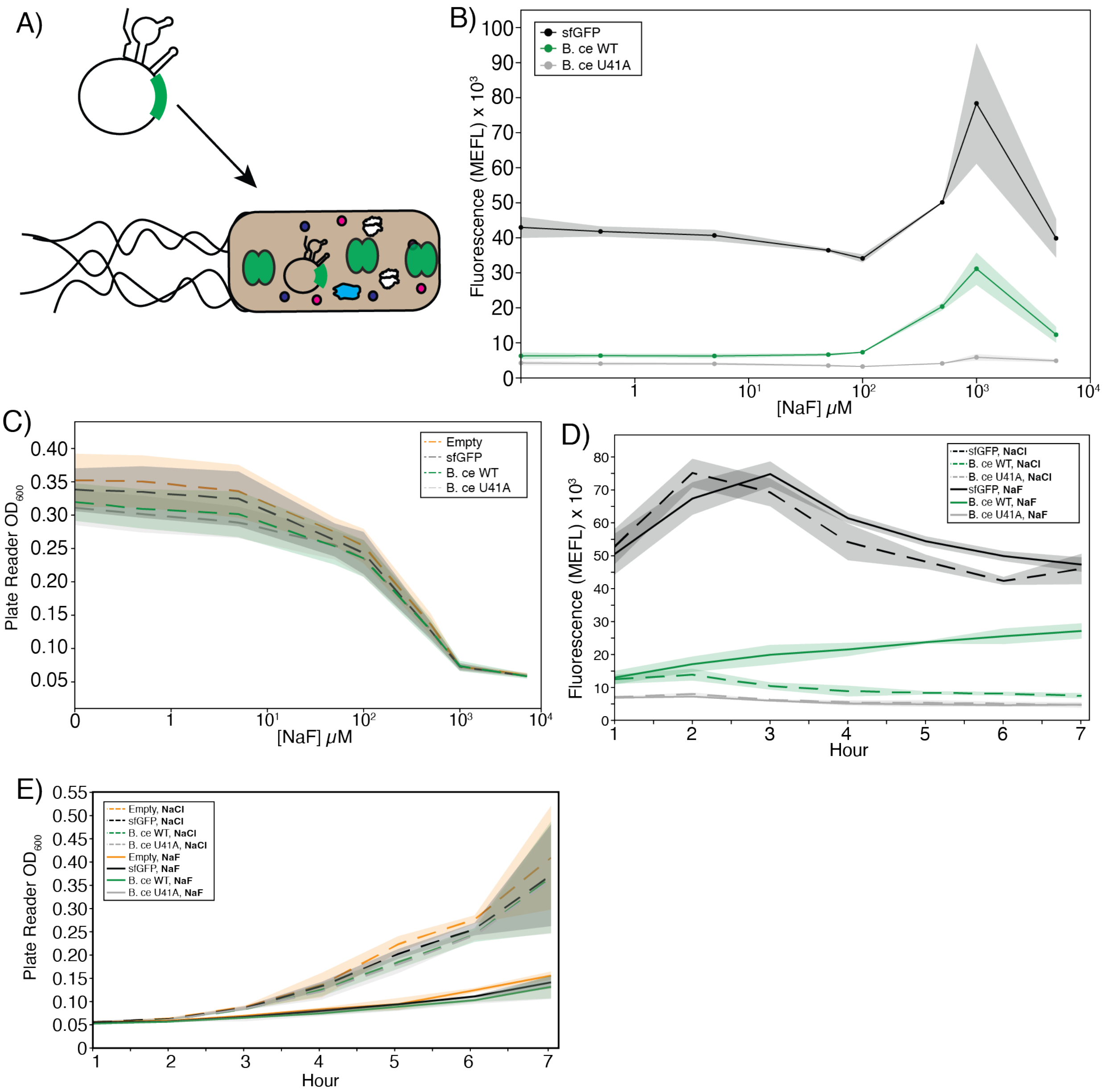


**Figure S1. Establishing a cellular gene expression assay for fluoride riboswitch function.** A) Cartoon representation of the riboswitch regulation of the reporter construct being expressed in cells. B) Cellular gene expression assay data collected through flow cytometry (single-cell quantification) after six hours of cell culture growth showing a dose response to externally supplied NaF using three constructs: constitutively expressed super-folded GFP (sfGFP), *B. ce* WT fluoride riboswitch, and the ligand unresponsive *B. ce* U41A fluoride riboswitch. Analysis of fold change shown in Figure 1D. C) $OD_{600}$ collected on a plate reader (bulk population quantification) of experiment from part B. D) Cellular gene expression assay time-course (single-cell flow cytometry) quantification collected for sfGFP, *B. ce* WT fluoride riboswitch, and the ligand unresponsive *B. ce* U41A fluoride riboswitch using either 500 μM NaCl or 500 μM NaF supplied to the media. E) $OD_{600}$ data collected on a plate reader (bulk population) of experiment from part D. Data in B-E represent averages (lines) and $\pm$ standard deviation (shading) for three biological replicates, each analyzed over three technical replicates (N=9). MEFL – molecules of equivalent fluorescein.

**Figure S2**

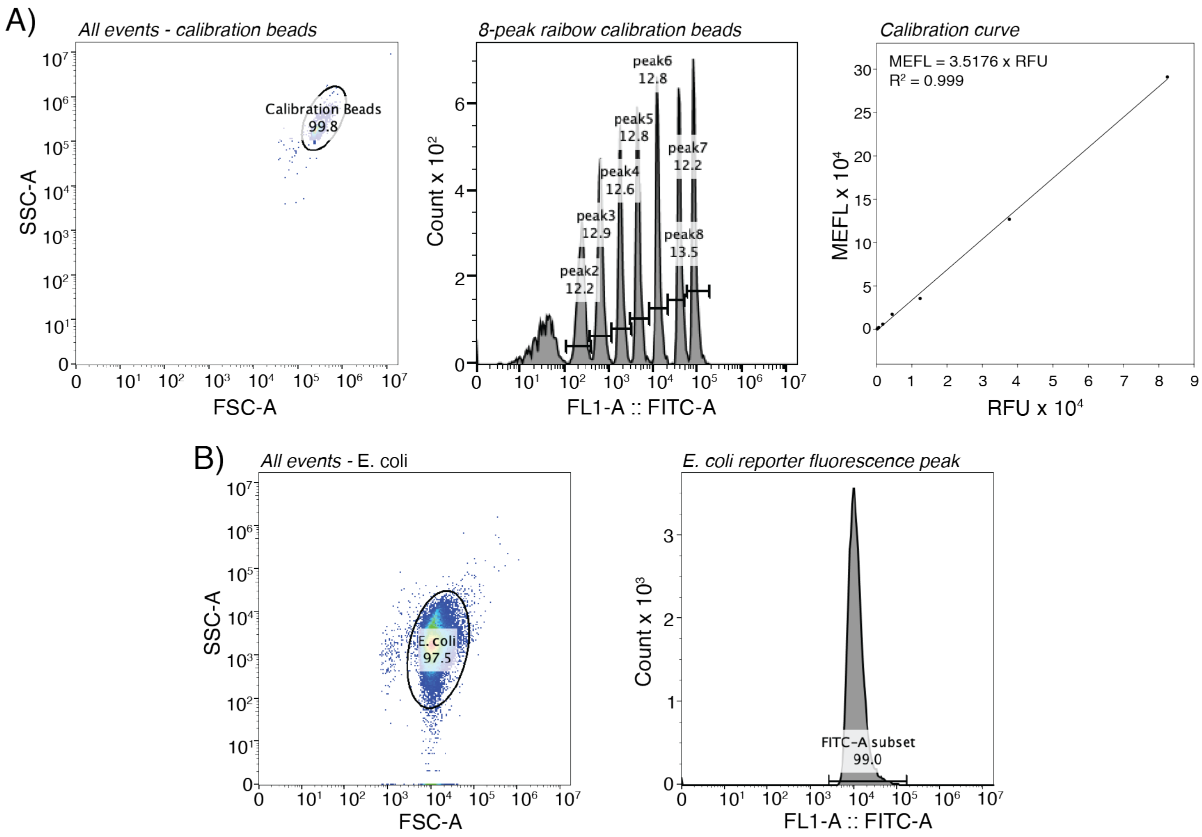


**Figure S2. Flow cytometry gating and calibration workflow.** A) Sphero$^{TM}$ calibration beads were used to convert fluorescence intensity to units of molecules of equivalent fluorescein (MEFL) (see **Methods**). Beads were run and first gated on SSC-A vs. FSC-A, resulting in eight different peaks being resolved in the FL1-A (FITC-A) channel. Gates were drawn around peaks 2-8 and the geometric mean of fluorescence was calculated for each gate to give an RFU value for that peak. Manufacture supplied MEFL values for each bead peak were plotted versus measured RFU, and a linear fit was performed (y intercept forced to 0) to establish a conversion factor between measured RFU and MEFL. B) Cellular populations were gated using an ellipsoid plotted around blank cells (not transformed with any plasmid) in the SSC-A vs. FSC-A channel. The same gate was applied to all cultures. Fluorescence populations were gated by sub-selecting the peak with the highest count. The same process was applied to all cultures. The geometric mean of the fluorescence was then calculated within this gate to obtain an RFU value, which was then converted to MEFL using the calibration curve. Fluorescence measurements were collected using BD CSampler software and peak averages and histograms were calculated using FlowJo 10.8.1.

### Figure S3

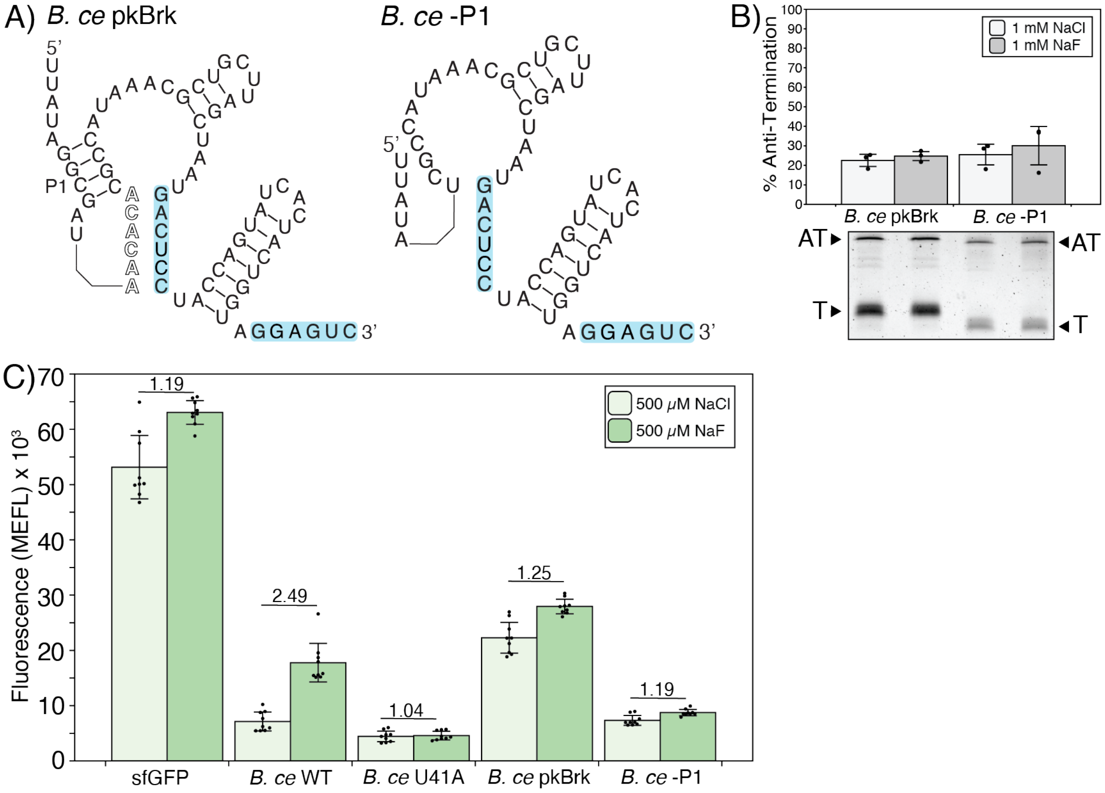


**Figure S3. Establishing a riboswitch terminator control.** A) Secondary structure schematics of two riboswitch controls that were designed to break (pkBrk) or remove (-P1) the pseudoknot, promoting terminator formation and thus favoring the riboswitch OFF state. B) Single-round *in vitro* transcription reactions for *B. ce* pkBrk and *B. ce* -P1 Stem constructs performed at 1 mM NaCl or NaF. A representative gel is shown under a plot of experimental replicates. C) The *B. ce* pkBrk and *B. ce* -P1 Stem constructs characterized in the cellular gene expression assay. Data in B represents three experimental replicates (N=3), with error bars representing standard deviation (raw gel images and replicates in Supplementary Data File 3). Bars in C represent average fluorescence for three biological replicates, each analyzed over three technical replicates (N=9) plotted as points, with error bars representing standard deviation, and numerical reporting of the fold change between the average from the 500 μM NaF and 500 μM NaCl conditions. MEFL – molecules of equivalent fluorescein.

**Figure S4**

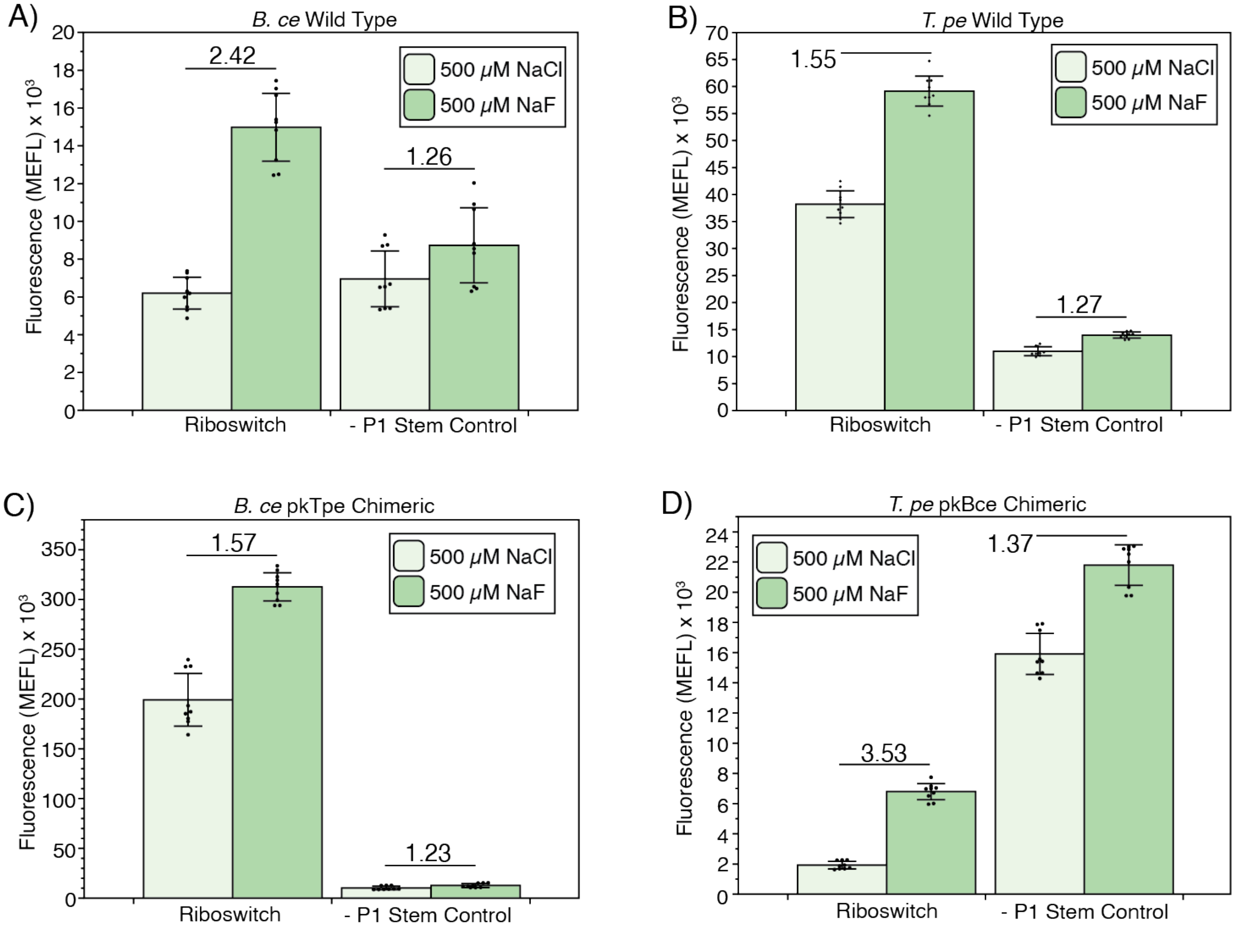


**Figure S4. Cellular gene expression controls for riboswitch constructs from Figure 2.** *B. ce* WT (A), *T. pe* WT (B), *B. ce* pkTpe (C), and *T. pe* pkBce (D) fluoride riboswitches from Figure 2 shown with their terminator control, where the P1 stem is removed (-P1). Bars represent average fluorescence for three biological replicates, each analyzed over three technical replicates (N=9) plotted as points, with error bars representing standard deviation, and numerical reporting of the fold change between the average from the 500 μM NaF and 500 μM NaCl conditions. MEFL – molecules of equivalent fluorescein.

**Figure S5**

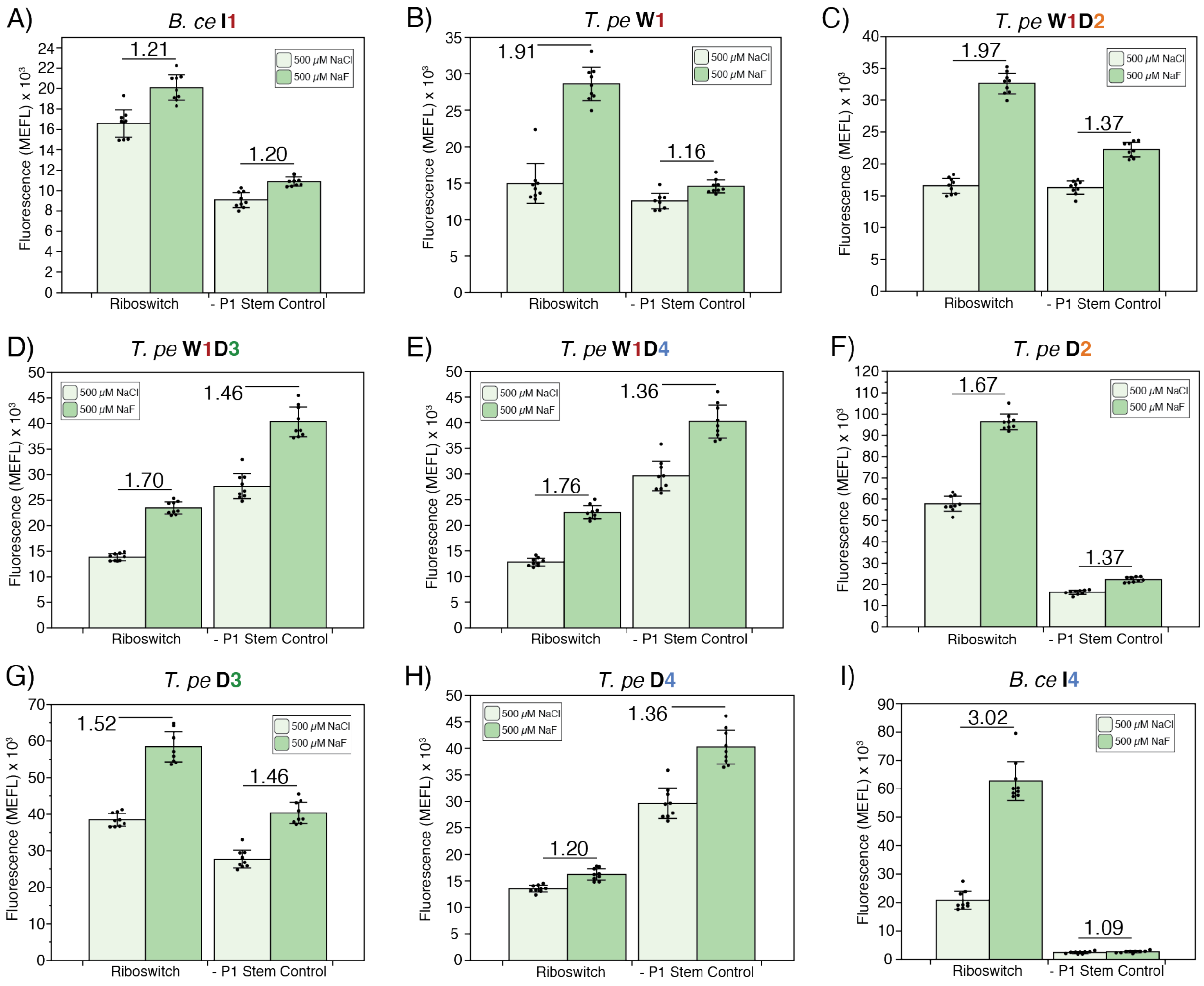


**Figure S5. Cellular gene expression controls for riboswitch constructs from Figure 3.** *B. ce* I1 (A), *T. pe* W1 (B), *T. pe* W1D2 (C), *T. pe* W1D3 (D), *T. pe* W1D4 (E), *T. pe* D2 (F), *T. pe* D3 (G), *T. pe* D4 (H), and *B. ce* I4 (I) fluoride riboswitches from Figure 3 shown with their terminator control, where the P1 stem is removed (-P1). Bars represent average fluorescence for three biological replicates, each analyzed over three technical replicates (N=9) plotted as points, with error bars representing standard deviation, and numerical reporting of the fold change between the average from the 500 μM NaF and 500 μM NaCl conditions. MEFL – molecules of equivalent fluorescein.

**Figure S6**

A)

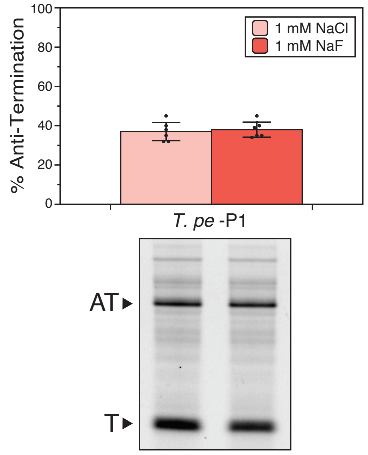


B)

| Values | ***T. pe* WT** | ***T. pe* W1** |
|---|---|---|
| Top | 94.21 | 91.15 |
| Bottom | 79.76 | 24.43 |
| Hillslope | 1.25 | 1.574 |
| AT50 | 1.523 | 1.921 |

**Figure S6. Control and equation variables for thermophilic transcription assay.** A) *In vitro* transcription reactions with *T. aq* RNAP at 65 °C for *T. pe* -P1 Stem variant in the presence of 1 mM NaCl or 1 mM NaF. A representative gel is shown. The Anti-Terminated (AT) (183 nts) and Terminated (T) (80 nts) bands are indicated. A representative gel is shown under a plot of experimental replicates. B) Table of values for the nonlinear regression curve fit:

$$(1)\ y = x^{Hillslope} \times \frac{(Top - Bottom)}{x^{Hillslope} + AT50^{Hillslope}} + Bottom$$

for the curves in Figure 4C as determined by GraphPad Prism 10. Data in A represents six experimental replicates (N=6), with error bars representing standard deviation (raw gel images and replicates in Supplementary Data File 3).

**Figure S7**

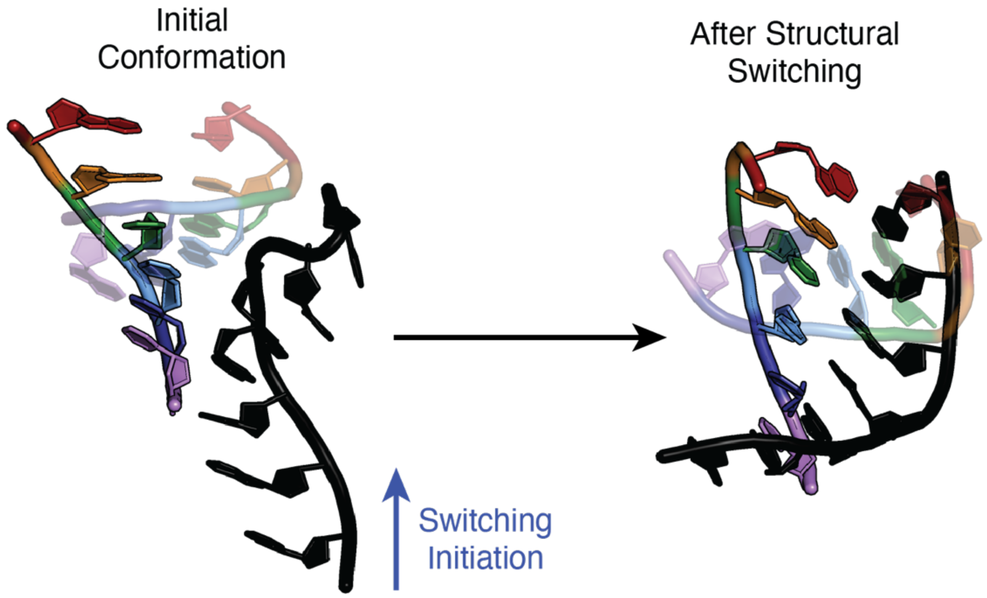


**Figure S7. Visualization of the 90° pitch of the end of the terminator from the pseudoknot.**

Pre- and post- structural switching through the pseudoknot (rainbow) by the terminator (black).

**Figure S8**

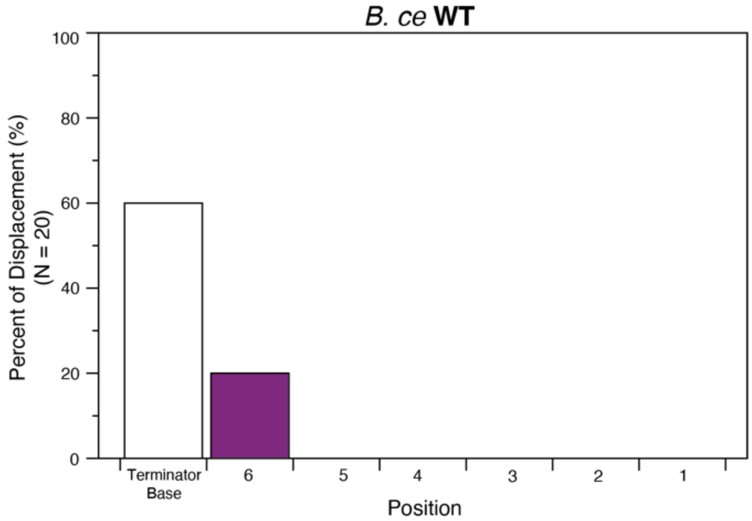


**Figure S8. Simulated structural switching in the absence of $Mg^{2+}$ for *B. ce* WT apo state.** Twenty replica structural switching simulations (N=20) of the apo states for the *B. ce* WT variant fluoride riboswitch were analyzed to quantify the percentage that the base of the terminator (U45-A64) formed and each pseudoknot position was displaced by expression platform folding at the end of the simulation. Bars are colored according to their position along the pseudoknot (Figure 1A) and the base of the terminator is in white. Note that even though all non-$Mg^{2+}$ *B. ce* WT simulations contained stronger biases than used in $Mg^{2+}$ containing simulations, no occurrences of complete strand exchange were observed indicating the essential role played by including diffuse $Mg^{2+}$ in the all-atom simulations.

**Supplementary Table S1**

| Primer | Sequence |
|---|---|
| A | GCTTCCGGCTTGATTCTAAAGATC |
| B | CGGACAGAAAATTTGTGCCC |
| C | CCGAATTCAAAAAGAGTATTGACTTAAAGTCTAACC |

**Supplementary Table S1.** Primer sequences used in PCR to generate dsDNA templates for single-round *in vitro* transcription.

**Supplementary Table S2**

| Atom 1 | Atom 2 | Function | Sigma (nm) | Epsilon (kJ/mol) |
|---|---|---|---|---|
| mMg | F | 1.000 | 0.3665 | 8.4679976 |

**Supplementary Table S2.** Non-bonded parameters for the site-bound $Mg_3F$, where the Sigma and Epsilon values are the modified Netz paramters.

**Supplementary Table S3**

| Atom 1 | Atom 2 | Strength (kJ/mol/nm$^2$) |
|---|---|---|
| mMg | F | 250 |
| mMg | O1P | 250 |
| mMg | O2P | 250 |
| mMg | O1P | 250 |

**Supplementary Table S3.** Distance restraints for site-bound $Mg_3F$.

**Supplementary Table S4**

***B. ce* WT & S1 Distance Restraints**

| Atom 1 | Atom 2 | Strength (kJ/mol/nm$^2$) |
|---|---|---|
| 1G H1 | 16C N3 | 250 |
| 4G H1 | 13C N3 | 250 |
| 23G H1 | 32C N3 | 250 |
| 44A N1 | 61U H3 | 250 |
| 50A N1 | 55U H3 | 250 |
| 37G H1 | 68C N3 | 175 |
| 38A N1 | 67U H3 | 175 |
| 39C N3 | 66G H1 | 175 |
| 40U H3 | 65A N1 | 175 |
| 41C N3 | 64G H1 | 175 |
| 42C N3 | 63G H1 | 175 |

*Bacillus cereus* (*B. ce*)

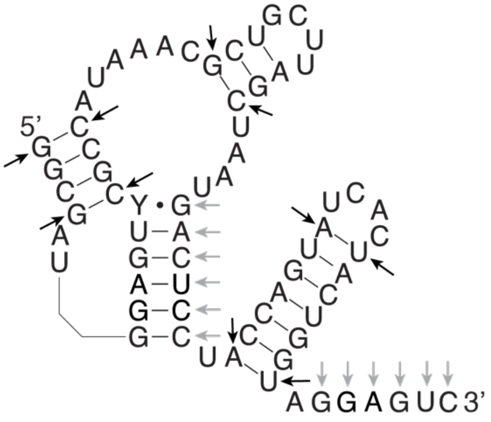

***T. pe* WT & W1 Distance Restraints**

| Atom 1 | Atom 2 | Strength (kJ/mol/nm$^2$) |
|---|---|---|
| 2G H1 | 17C N3 | 250 |
| 5G H1 | 14C N3 | 250 |
| 24G H1 | 37C N3 | 250 |
| 49A N1 | 63U H3 | 250 |
| 53G H1 | 59U O2 | 250 |
| 42G H1 | 70C N3 | 175 |
| 43G H1 | 69C N3 | 175 |
| 44C N3 | 68G H1 | 175 |
| 45C N3 | 67G H1 | 175 |
| 46U H3 | 66A N1 | 175 |
| 47C N3 | 65G H1 | 175 |

*Thermotoga petrophila* (*T. pe*)

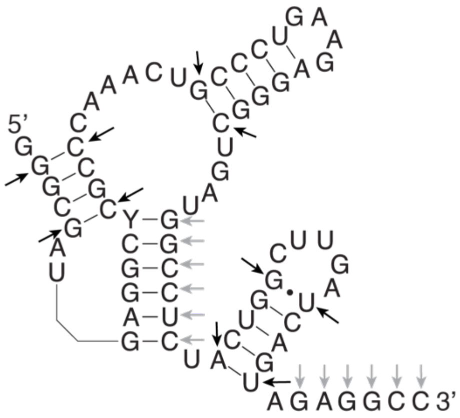

**Supplementary Table S4.** Distance restraints between pairs of atoms in the *B. ce* WT/ S1 and *T. pe* WT/ W1 variants. 2D schematics are shown on the right with black and grey arrows pointing to locations with a restraint of 250 kJ/mol/nm$^2$ and 175 kJ/mol/nm$^2$, respectively. The distance restraints are enforced, i.e. the designated energetic penalty is applied, if the distance between the atom pair is > 4 Å or < 1.6 Å as the potential for these restraints is flat-bottomed.

**Supplementary Table S5**

| Run No. | Model | Purpose | Length of Simulation (ns) |
|---|---|---|---|
| 1 | Apo State WT & Mutant Homology Models (*B. ce* & *T. pe*) | Equilibration | 11 (x4) |
| 2 | Holo State WT & Mutant Homology Models (*B. ce* & *T. pe*) | Equilibration | 11 (x4) |
| 3 | Equilibrated Apo State *B. ce* WT Model | Biased Strand Displacement | 40 (x20) |
| 4 | Equilibrated Apo State *B. ce* Mutant Model | Biased Strand Displacement | 40 (x20) |
| 5 | Equilibrated Holo State *B. ce* WT Model | Biased Strand Displacement | 40 (x20) |
| 6 | Equilibrated Holo State *B.ce* Mutant Model | Biased Strand Displacement | 40 (x20) |
| 7 | Equilibrated Apo State *T.pe* WT Model | Biased Strand Displacement | 40 (x20) |
| 8 | Equilibrated Apo State *T.pe* Mutant Model | Biased Strand Displacement | 40 (x20) |
| 9 | Equilibrated Holo State *T.pe* WT Model | Biased Strand Displacement | 40 (x20) |
| 10 | Equilibrated Holo State *T.pe* Mutant Model | Biased Strand Displacement | 40 (x20) |
| Total | | | 6,488 |

**Supplementary Table S5.** Summary of molecular dynamics simulations performed.

**Supplementary Movie 1. All-atom molecular dynamics simulations of apo state of the *B. ce* wild type fluoride riboswitch.** The video shows a representative molecular dynamics (MD) trajectory of structural switching in the *B. ce* WT apo model. The simulation equilibrates the 68 nt model then applies flat-bottom, harmonic distance restraints. The invading strand is colored in black, and the pseudoknot is colored according to the scheme shown in Figure 1A.

**Supplementary Movie 2. All-atom molecular dynamics simulations of holo state of the *B. ce* wild type fluoride riboswitch.** The video shows a representative molecular dynamics (MD) trajectory of structural switching in the *B. ce* WT holo model, which includes a $Mg_3F$ ligand with pre-formed, non-dissociating ion-RNA contacts. The simulation equilibrates the 68 nt model then applies flat-bottom, harmonic distance restraints. The invading strand is colored in black, and the pseudoknot is colored according to the scheme shown in Figure 1A.

**Supplementary Movie 3. All-atom molecular dynamics simulations of apo state of the *B. ce* I1 variant fluoride riboswitch.** The video shows a representative molecular dynamics (MD) trajectory of structural switching in the *B. ce* I1 (S1 naming in movie) variant apo model. The simulation equilibrates the 68 nt model then applies flat-bottom, harmonic distance restraints. The invading strand is colored in black, and the pseudoknot is colored according to the scheme shown in Figure 1A.

**Supplementary Data File 1.** Expression construct sequences used in this study.

**Supplementary Data File 2.** All source data of the calibrated flow cytometry data and band intensity analysis.

**Supplementary Data File 3.** Raw and replicated gel images and text descriptions.
