## Supplementary Data File 3 for "The Effect of Pseudoknot Base Pairing on Cotranscriptional Structural Switching of the Fluoride Riboswitch"

**Document of Raw Gel Images For:**

Figure 1C and S3b – Raw Gel Replicate 1

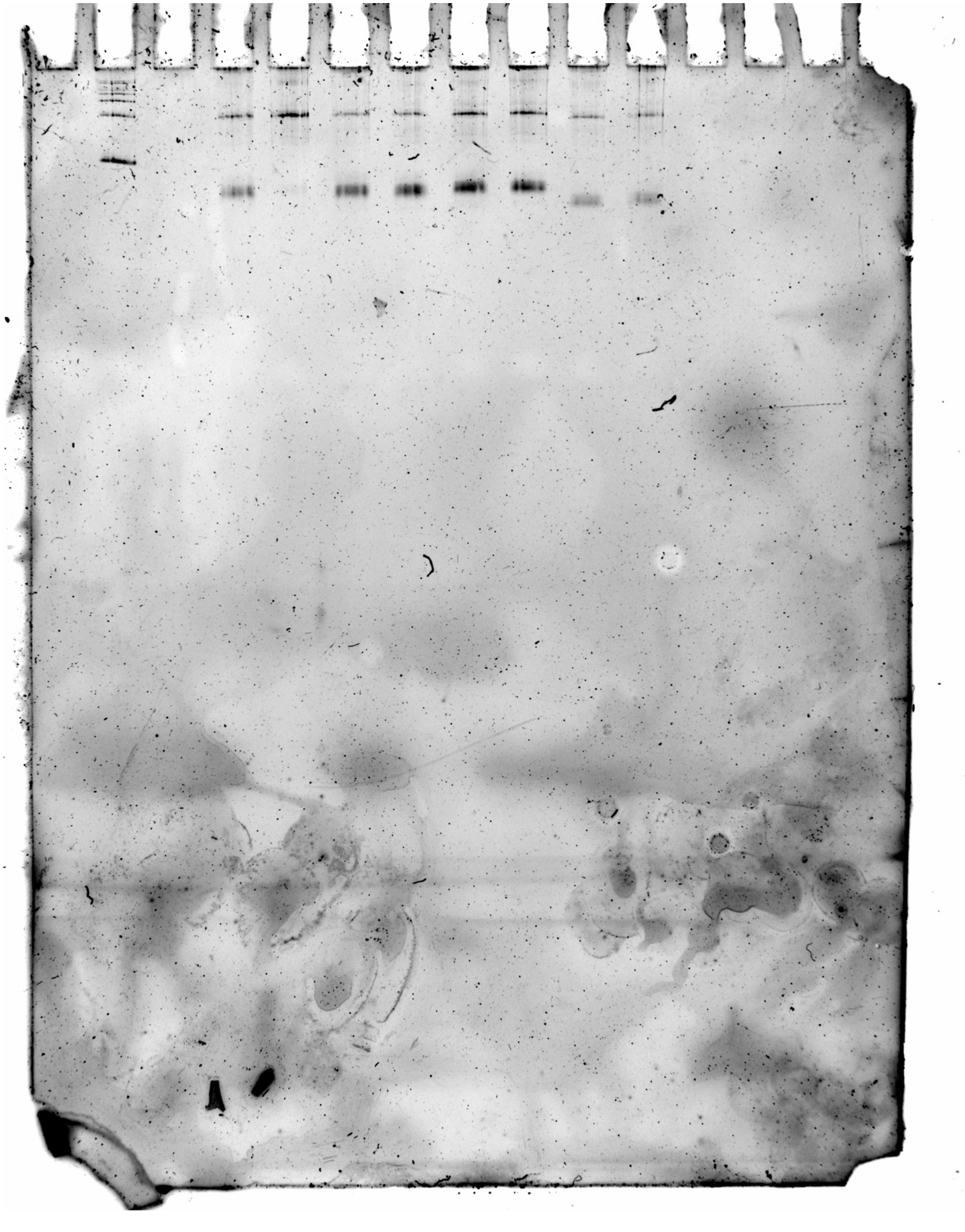

Figure 1C and S3b – Raw Gel Replicate 2

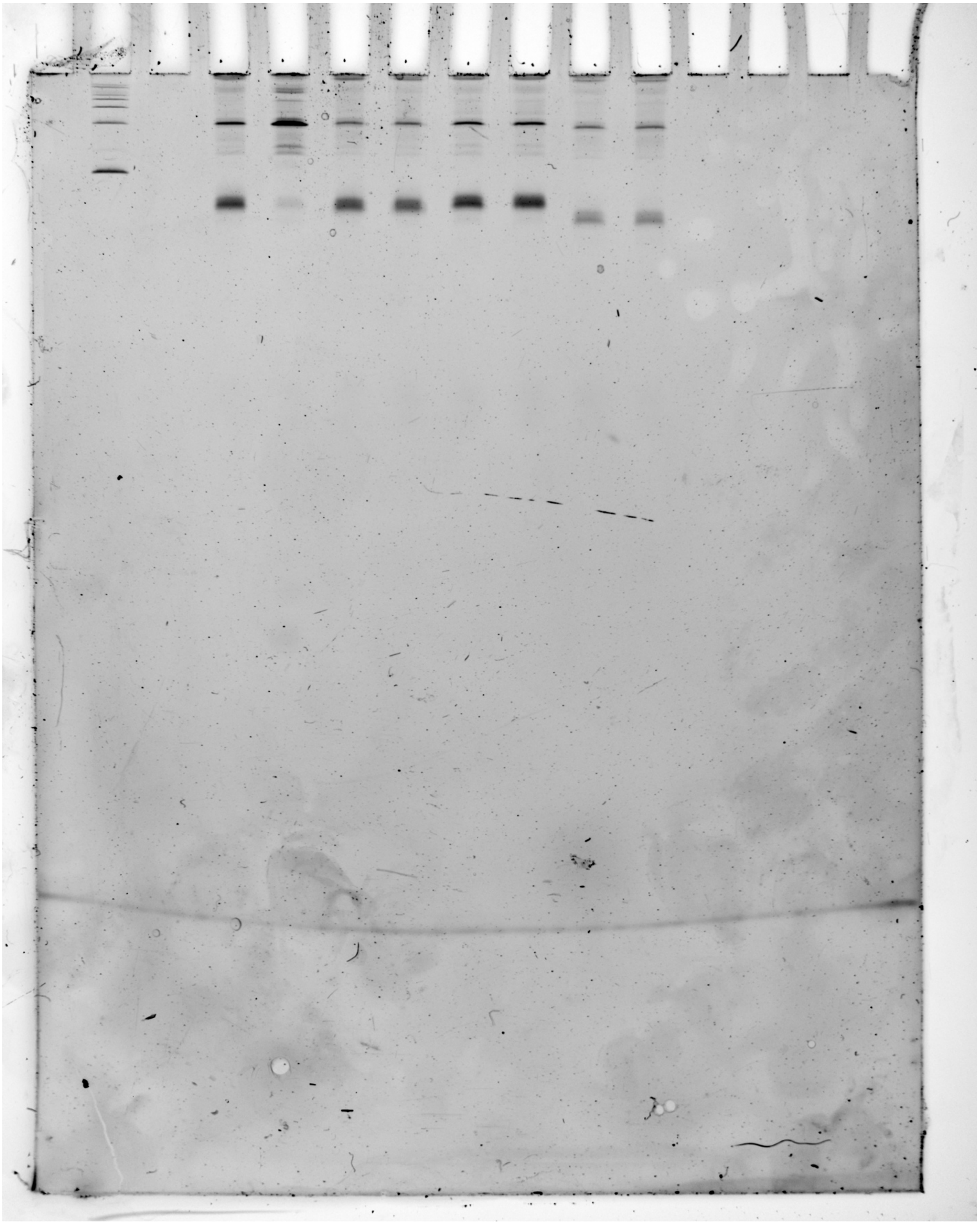

Figure 1C, 3G, and S3b – Raw Gel Replicate 3

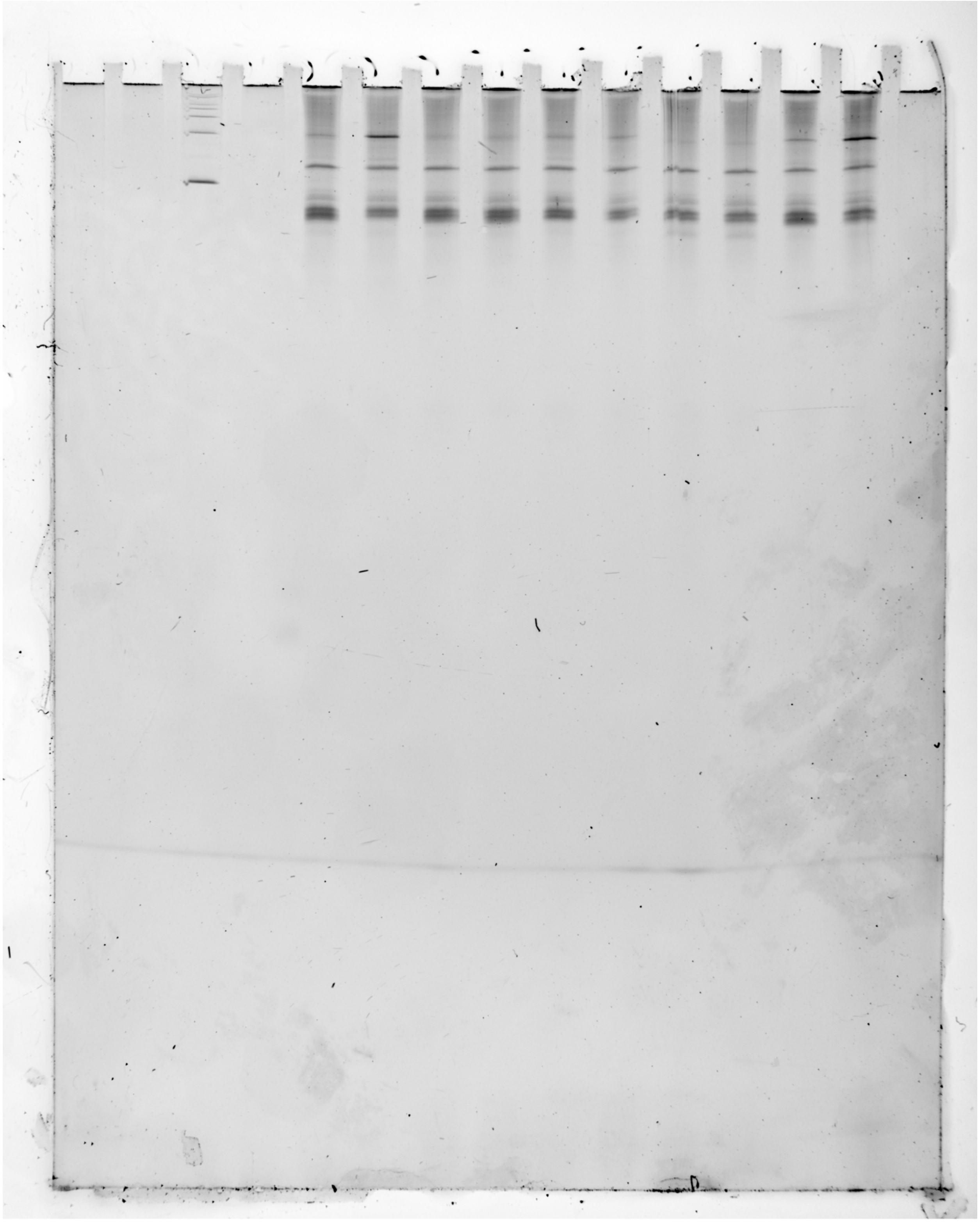

Figure 1C, 3G, and S3b – Notes

1c_S3b_Rep1, 1c_S3b_Rep2, 1c_3g_S3b_Rep3 all show the uncrossed, unprocessed urea-PAGE gel image of the data shown in Fig. 1c and Fig. S3b (and Fig. 3g), including the other two replicates not explicitly shown in the panel. 1c_S3b_Rep2 is the uncrossed, unprocessed image shown in the manuscript for both figures. Lane 1-3 and 12-15 (for 1c_3g_S3b_Rep3: 1-4 and 15) are not shown in the manuscript.

From left to right:
2. ssRNA Ladder (100, 200, 300, 400, 500, 750, and 1000 bases) (Invitrogen, cat. no. AM7145)
4. In vitro transcript products generated from E. coli RNAp during a single-round of transcription with 1 mM NaCl, DNA template = B. ce Fluoride Riboswitch WT
5. In vitro transcript products generated from E. coli RNAp during a single-round of transcription with 1 mM NaF, DNA template = B. ce Fluoride Riboswitch WT
6. In vitro transcript products generated from E. coli RNAp during a single-round of transcription with 1 mM NaCl, DNA template = B. ce Fluoride Riboswitch U41A
7. In vitro transcript products generated from E. coli RNAp during a single-round of transcription with 1 mM NaF, DNA template = B. ce Fluoride Riboswitch U41A
8. In vitro transcript products generated from E. coli RNAp during a single-round of transcription with 1 mM NaCl, DNA template = B. ce Fluoride Riboswitch pkBrk
9. In vitro transcript products generated from E. coli RNAp during a single-round of transcription with 1 mM NaF, DNA template = B. ce Fluoride Riboswitch pkBrk
10. In vitro transcript products generated from E. coli RNAp during a single-round of transcription with 1 mM NaCl, DNA template = B. ce Fluoride Riboswitch -P1
11. In vitro transcript products generated from E. coli RNAp during a single-round of transcription with 1 mM NaF, DNA template = B. ce Fluoride Riboswitch -P1
(For 1c_3g_S3b_Rep3)
13. In vitro transcript products generated from E. coli RNAp during a single-round of transcription with 1 mM NaCl, DNA template = B. ce Fluoride Riboswitch I4
14. In vitro transcript products generated from E. coli RNAp during a single-round of transcription with 1 mM NaF, DNA template = B. ce Fluoride Riboswitch I4

Figure 2D – Raw Gel Replicate 1

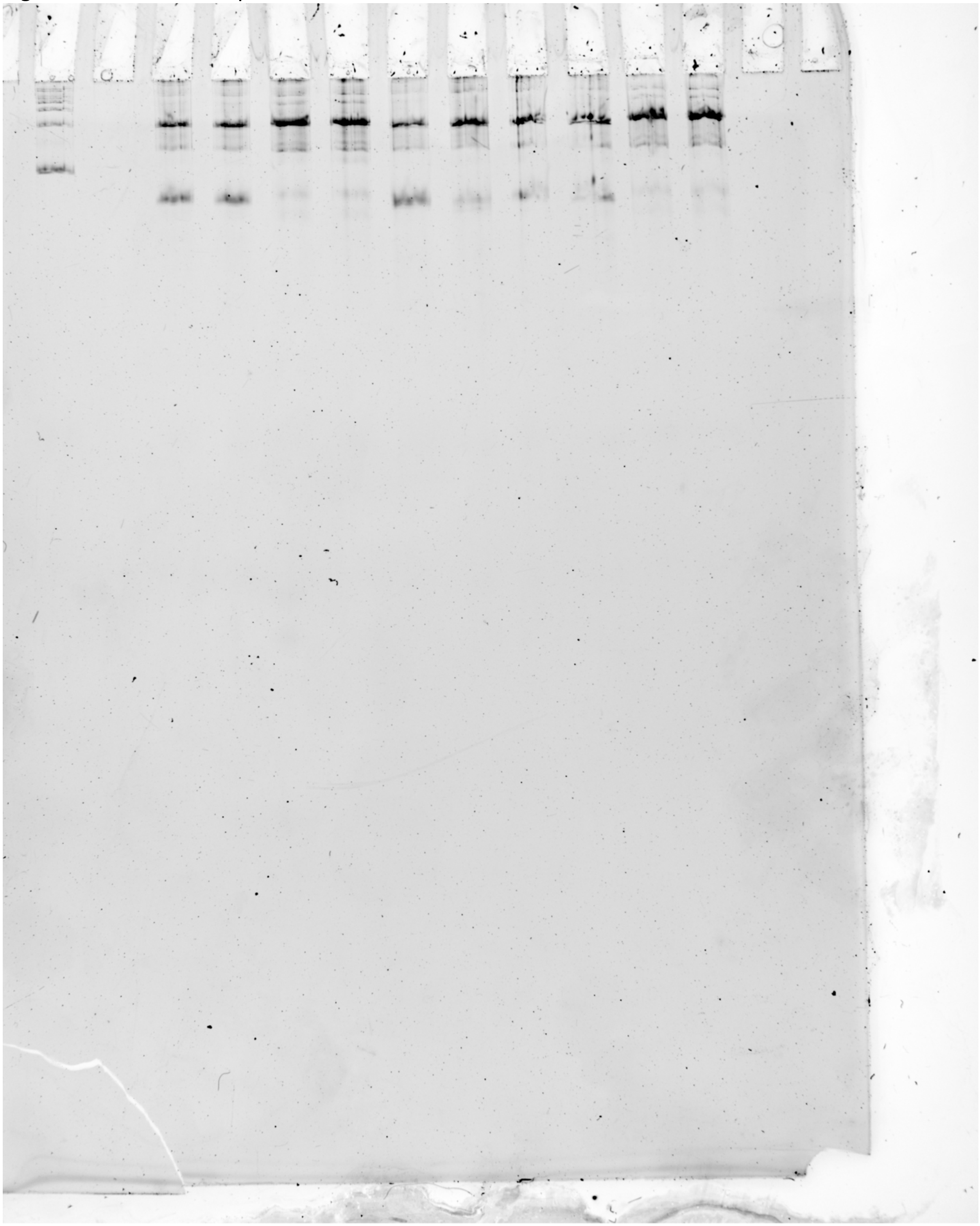

Figure 2D – Raw Gel Replicate 2

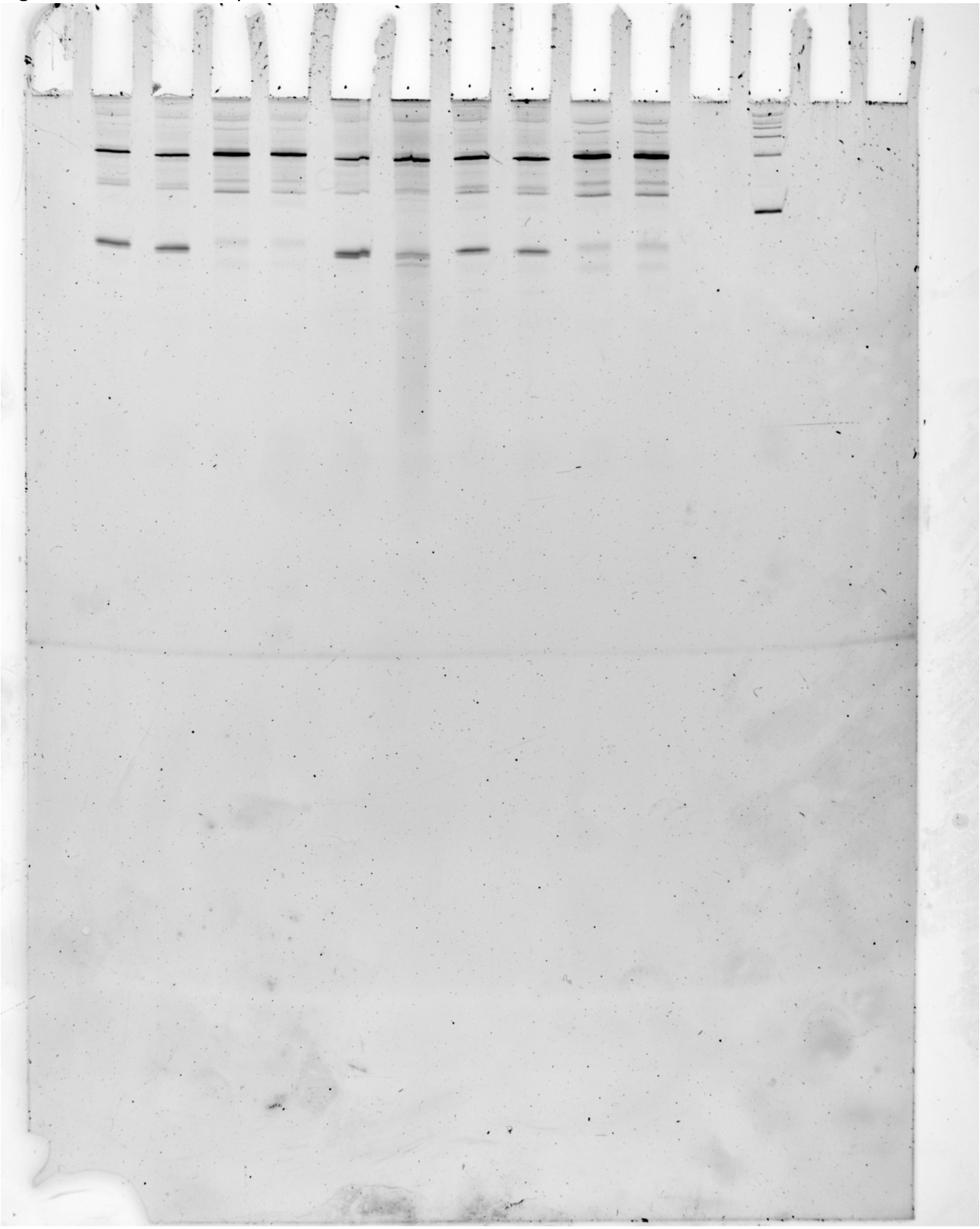

Figure 2D – Raw Gel Replicate 3

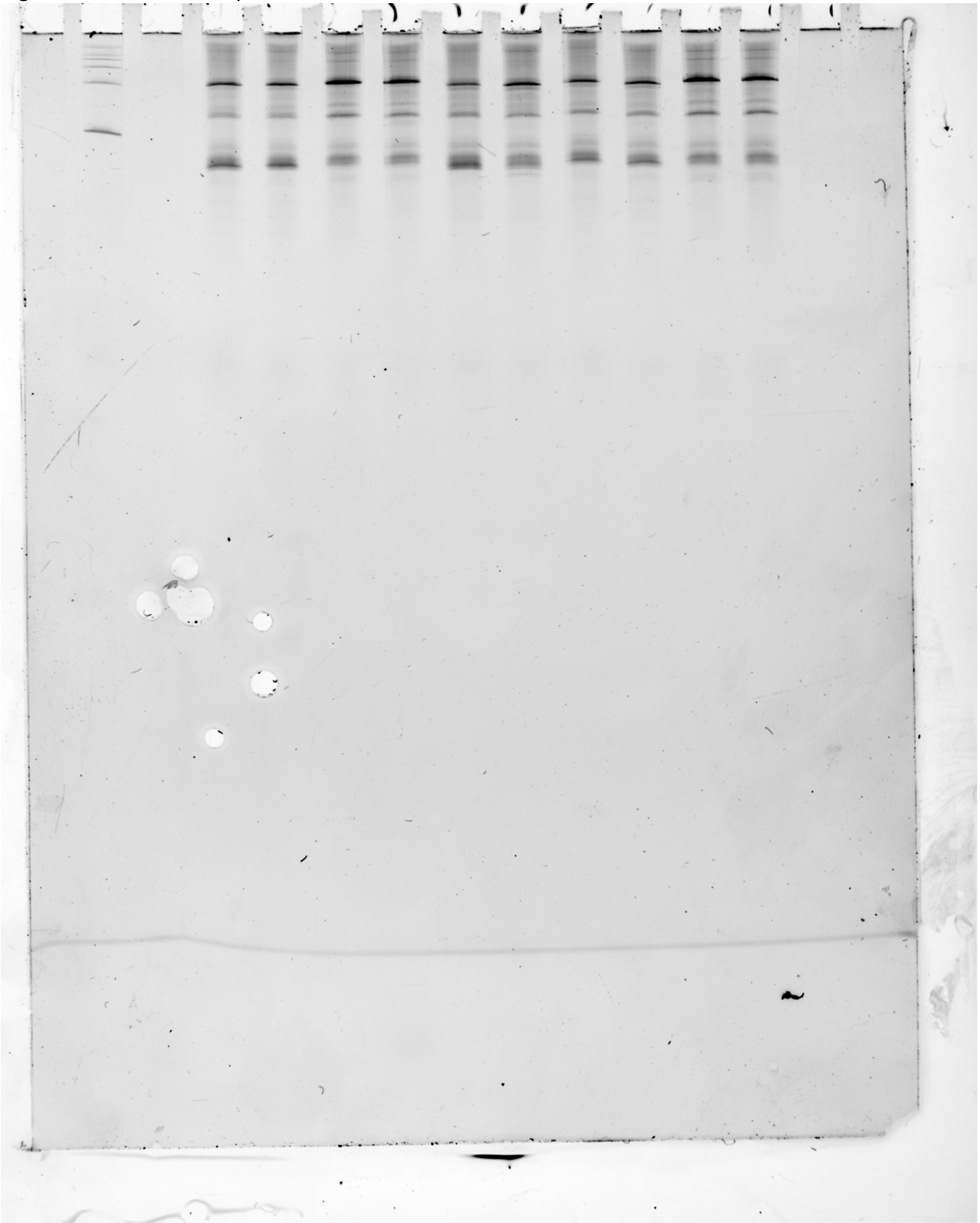

Figure 2D – Notes

2d_Rep1, 2d_Rep2, and 2d_Rep3 all show the uncrossed, unprocessed urea-PAGE gel image of the data shown in Fig. 2d, including the other two replicates not explicitly shown in the panel. 2d_Rep2 is the uncrossed, unprocessed image shown in the manuscript for both figures. Lanes 1 and 8-15 are not shown in the manuscript.

From left to right (2d_Rep2):
2. In vitro transcript products generated from E. coli RNAp during a single-round of transcription with 1 mM NaCl, DNA template = T. pe Fluoride Riboswitch WT
3. In vitro transcript products generated from E. coli RNAp during a single-round of transcription with 1 mM NaF, DNA template = T. pe Fluoride Riboswitch WT
4. In vitro transcript products generated from E. coli RNAp during a single-round of transcription with 1 mM NaCl, DNA template = B. ce Fluoride Riboswitch pkTpe
5. In vitro transcript products generated from E. coli RNAp during a single-round of transcription with 1 mM NaF, DNA template = B. ce Fluoride Riboswitch pkTpe
6. In vitro transcript products generated from E. coli RNAp during a single-round of transcription with 1 mM NaCl, DNA template = T. pe Fluoride Riboswitch pkBce
7. In vitro transcript products generated from E. coli RNAp during a single-round of transcription with 1 mM NaF, DNA template = T. pe Fluoride Riboswitch pkBce
8. In vitro transcript products generated from E. coli RNAp during a single-round of transcription with 1 mM NaCl, DNA template = T. pe Fluoride Riboswitch U44A
9. In vitro transcript products generated from E. coli RNAp during a single-round of transcription with 1 mM NaF, DNA template = T. pe Fluoride Riboswitch U44A
10. In vitro transcript products generated from E. coli RNAp during a single-round of transcription with 1 mM NaCl, DNA template = B. ce Fluoride Riboswitch pkTpe U41A
11. In vitro transcript products generated from E. coli RNAp during a single-round of transcription with 1 mM NaCl, DNA template = B. ce Fluoride Riboswitch pkTpe U41A
13. ssRNA Ladder (100, 200, 300, 400, 500, 750, and 1000 bases) (Invitrogen, cat. no. AM7145)

From left to right (2d_Rep1 and 2d_Rep3):
2. ssRNA Ladder (100, 200, 300, 400, 500, 750, and 1000 bases) (Invitrogen, cat. no. AM7145)
4. In vitro transcript products generated from E. coli RNAp during a single-round of transcription with 1 mM NaCl, DNA template = T. pe Fluoride Riboswitch WT
5. In vitro transcript products generated from E. coli RNAp during a single-round of transcription with 1 mM NaF, DNA template = T. pe Fluoride Riboswitch WT
6. In vitro transcript products generated from E. coli RNAp during a single-round of transcription with 1 mM NaCl, DNA template = B. ce Fluoride Riboswitch pkTpe
7. In vitro transcript products generated from E. coli RNAp during a single-round of transcription with 1 mM NaF, DNA template = B. ce Fluoride Riboswitch pkTpe
8. In vitro transcript products generated from E. coli RNAp during a single-round of transcription with 1 mM NaCl, DNA template = T. pe Fluoride Riboswitch pkBce

9. In vitro transcript products generated from E. coli RNAp during a single-round of transcription with 1 mM NaF, DNA template = T. pe Fluoride Riboswitch pkBce
10. In vitro transcript products generated from E. coli RNAp during a single-round of transcription with 1 mM NaCl, DNA template = T. pe Fluoride Riboswitch U44A
11. In vitro transcript products generated from E. coli RNAp during a single-round of transcription with 1 mM NaF, DNA template = T. pe Fluoride Riboswitch U44A
12. In vitro transcript products generated from E. coli RNAp during a single-round of transcription with 1 mM NaCl, DNA template = B. ce Fluoride Riboswitch pkTpe U41A
13. In vitro transcript products generated from E. coli RNAp during a single-round of transcription with 1 mM NaCl, DNA template = B. ce Fluoride Riboswitch pkTpe U41A

Figure 3C and 3G – Raw Gel Replicate 1

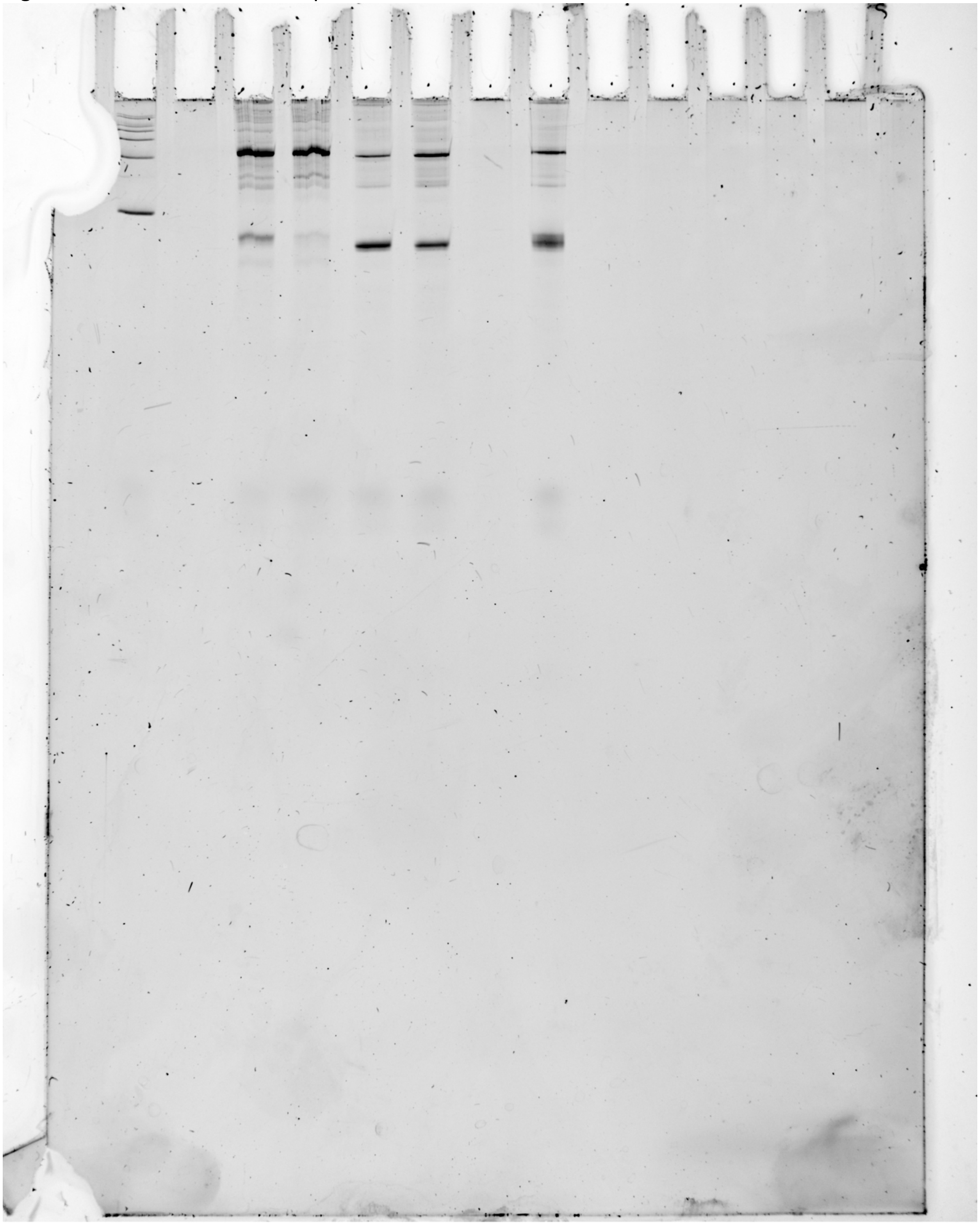

Figure 3C and 3G – Raw Gel Replicate 2

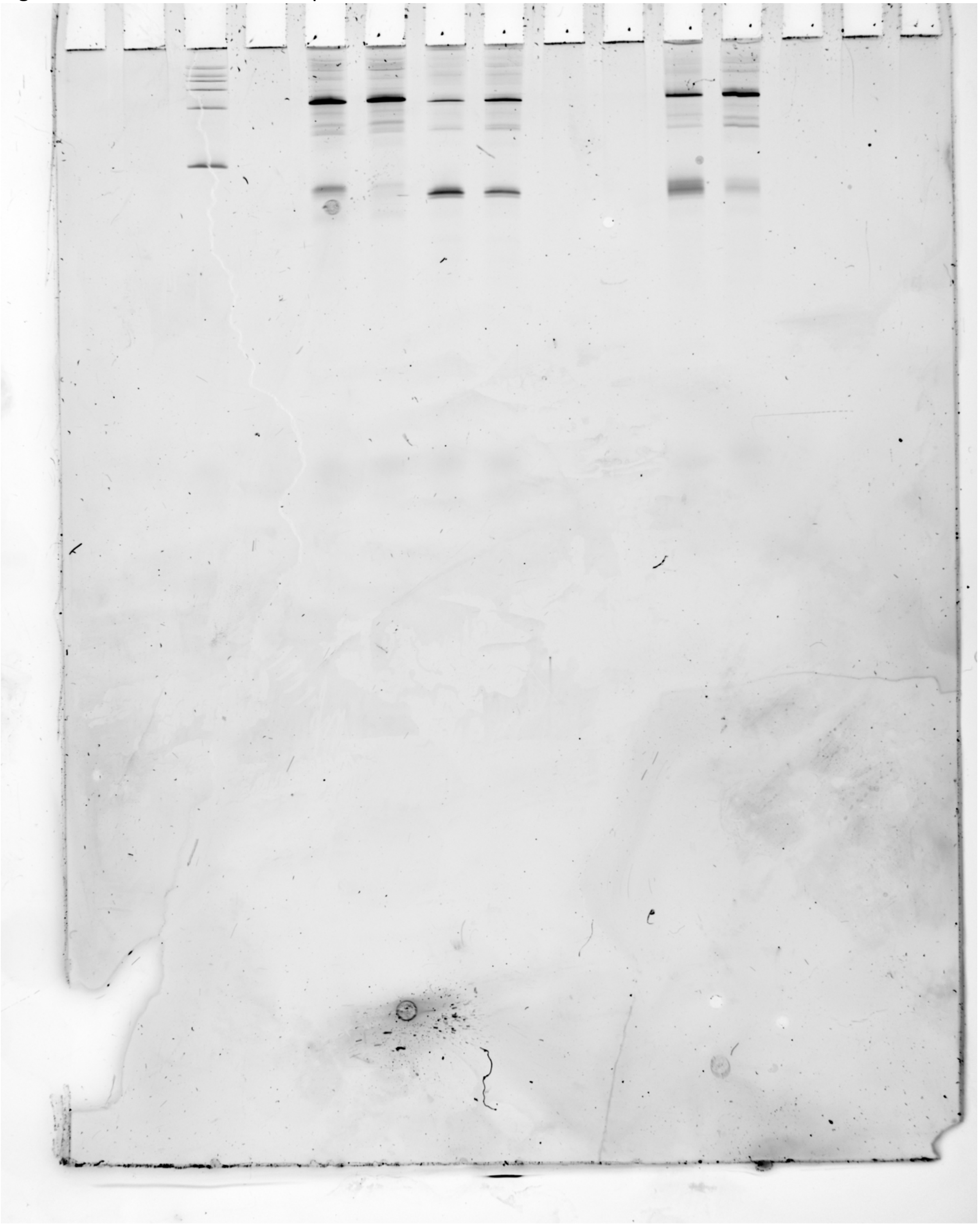

Figure 3C and 3G – Raw Gel Replicate 3

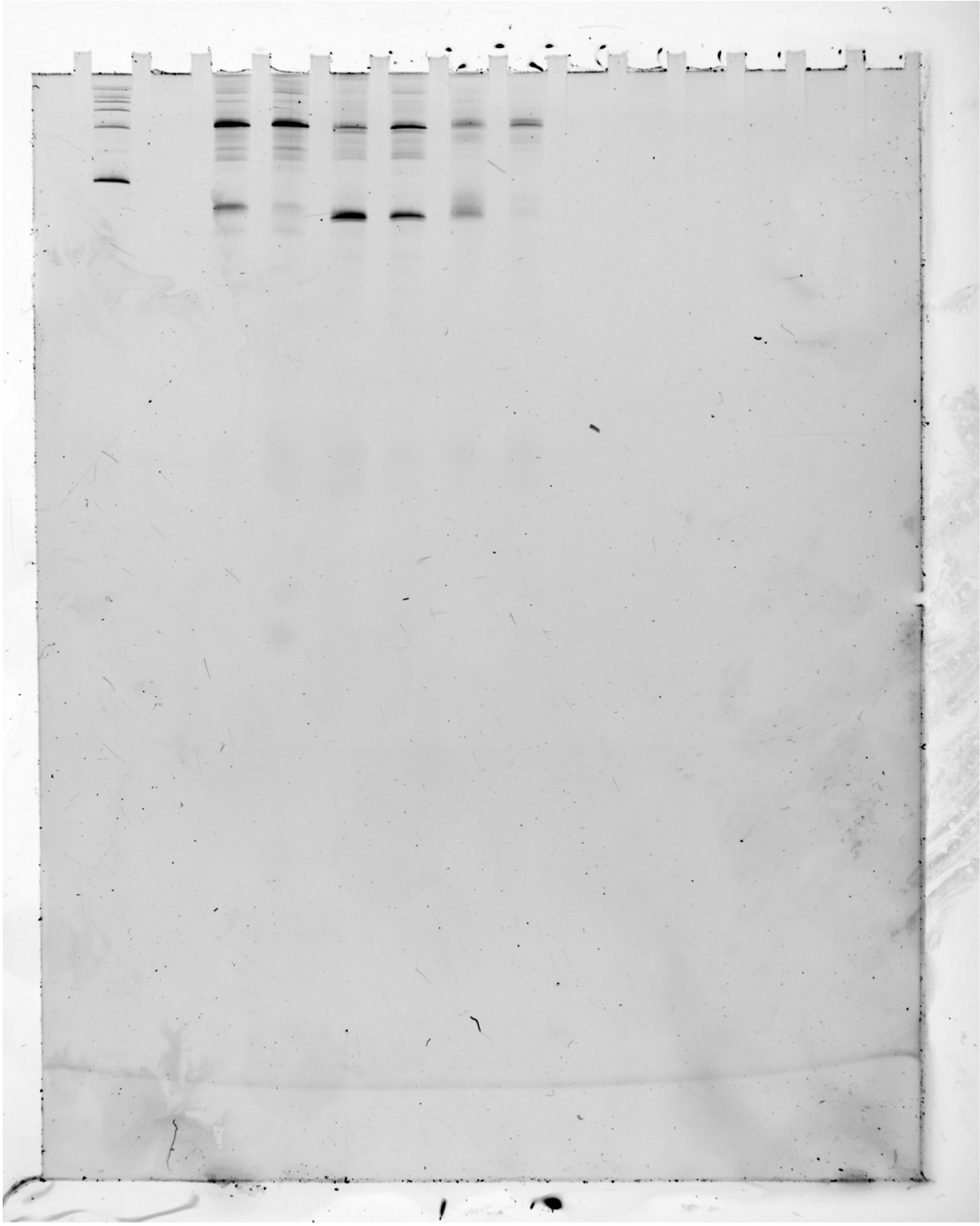

Figure 3C and 3G – Notes

3c_3g_Rep1, 3c_3g_Rep2, and 3c_3g_Rep3 all show the uncrossed, unprocessed urea-PAGE gel image of the data shown in Fig. 3c and Fig. 3g, including the other two replicates not explicitly shown in the panel. 3c_3g_Rep2 is the uncrossed, unprocessed image shown in the manuscript for both figures. Lanes 1-4, 9-10, and 13-15 are not shown in the manuscript.

From left to right (For 3c_3g_Rep2):
3. ssRNA Ladder (100, 200, 300, 400, 500, 750, and 1000 bases) (Invitrogen, cat. no. AM7145)
5. In vitro transcript products generated from E. coli RNAp during a single-round of transcription with 1 mM NaCl, DNA template = B. ce Fluoride Riboswitch S1
6. In vitro transcript products generated from E. coli RNAp during a single-round of transcription with 1 mM NaF, DNA template = B. ce Fluoride Riboswitch S1
7. In vitro transcript products generated from E. coli RNAp during a single-round of transcription with 1 mM NaCl, DNA template = T. pe Fluoride Riboswitch W1
8. In vitro transcript products generated from E. coli RNAp during a single-round of transcription with 1 mM NaF, DNA template = T. pe Fluoride Riboswitch W1
11. In vitro transcript products generated from E. coli RNAp during a single-round of transcription with 1 mM NaCl, DNA template = B. ce Fluoride Riboswitch S4
12. In vitro transcript products generated from E. coli RNAp during a single-round of transcription with 1 mM NaF, DNA template = B. ce Fluoride Riboswitch S4

From left to right (For 3c_3g_Rep1 and 3c_3g_Rep3):
2. ssRNA Ladder (100, 200, 300, 400, 500, 750, and 1000 bases) (Invitrogen, cat. no. AM7145)
4. In vitro transcript products generated from E. coli RNAp during a single-round of transcription with 1 mM NaCl, DNA template = B. ce Fluoride Riboswitch S1
5. In vitro transcript products generated from E. coli RNAp during a single-round of transcription with 1 mM NaF, DNA template = B. ce Fluoride Riboswitch S1
6. In vitro transcript products generated from E. coli RNAp during a single-round of transcription with 1 mM NaCl, DNA template = T. pe Fluoride Riboswitch W1
7. In vitro transcript products generated from E. coli RNAp during a single-round of transcription with 1 mM NaF, DNA template = T. pe Fluoride Riboswitch W1
8. In vitro transcript products generated from E. coli RNAp during a single-round of transcription with 1 mM NaCl, DNA template = B. ce Fluoride Riboswitch S4 (3c_3g_Rep1.tif: SAMPLE LOST DURING ANALYSIS PREPARATION, see 1c_3g_S3b_Rep3.tif for repeat)
9. In vitro transcript products generated from E. coli RNAp during a single-round of transcription with 1 mM NaF, DNA template = B. ce Fluoride Riboswitch S4 (3c_3g_Rep1.tif: see 1c_3g_S3b_Rep3.tif for repeat to match Lane 8)

Figure 3E – Raw Gel Replicate 1

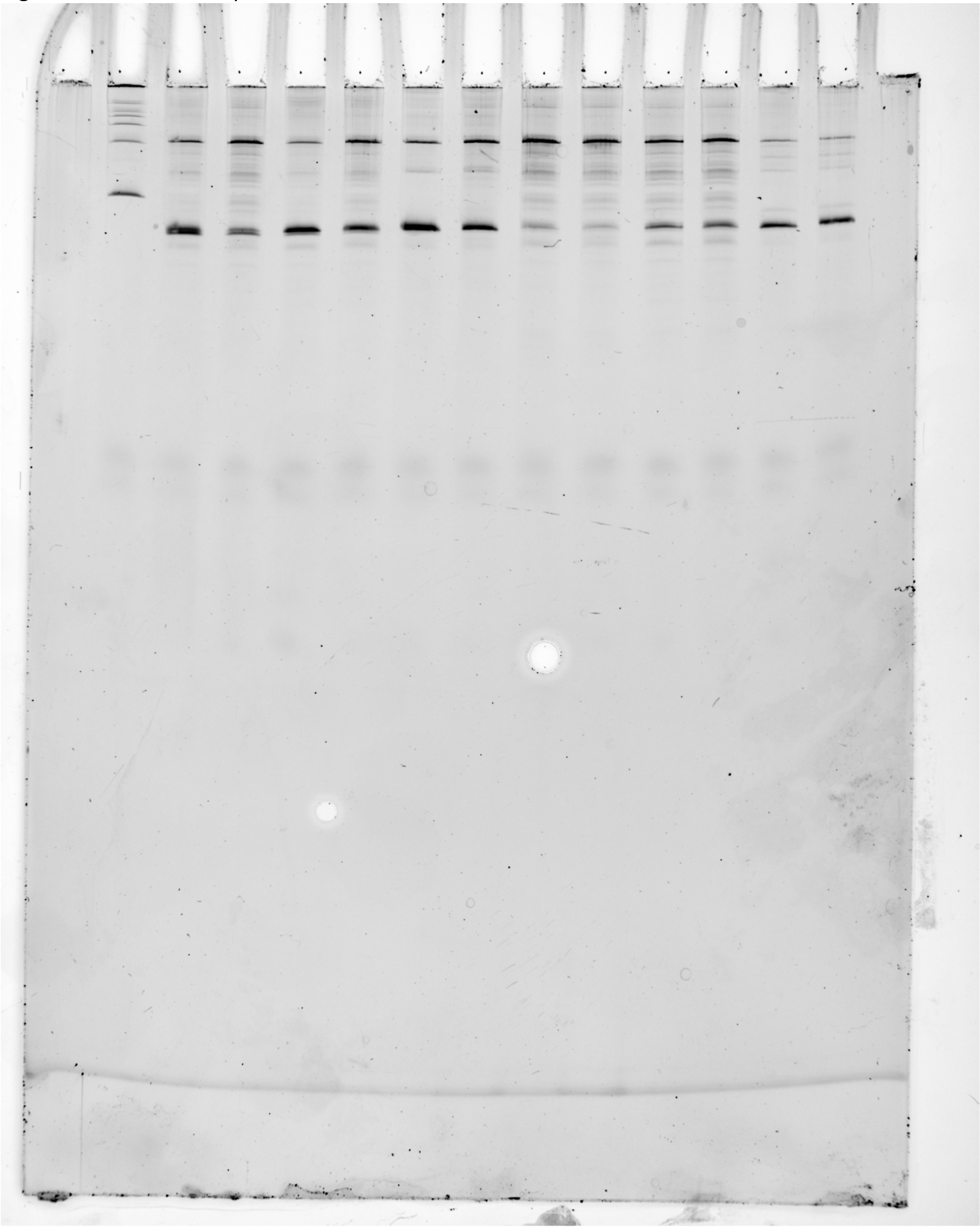

Figure 3E – Raw Gel Replicate 2

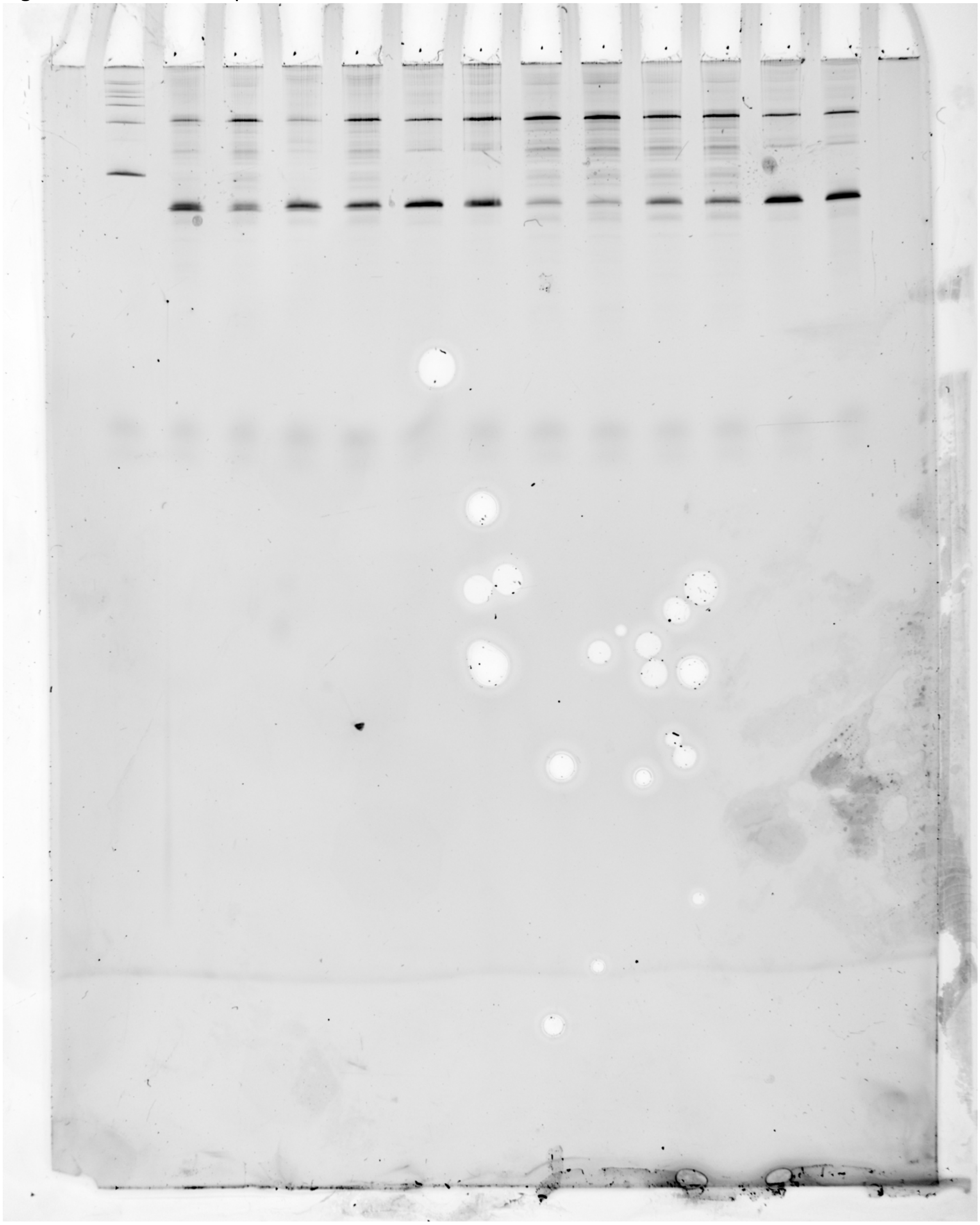

Figure 3E – Raw Gel Replicate 3

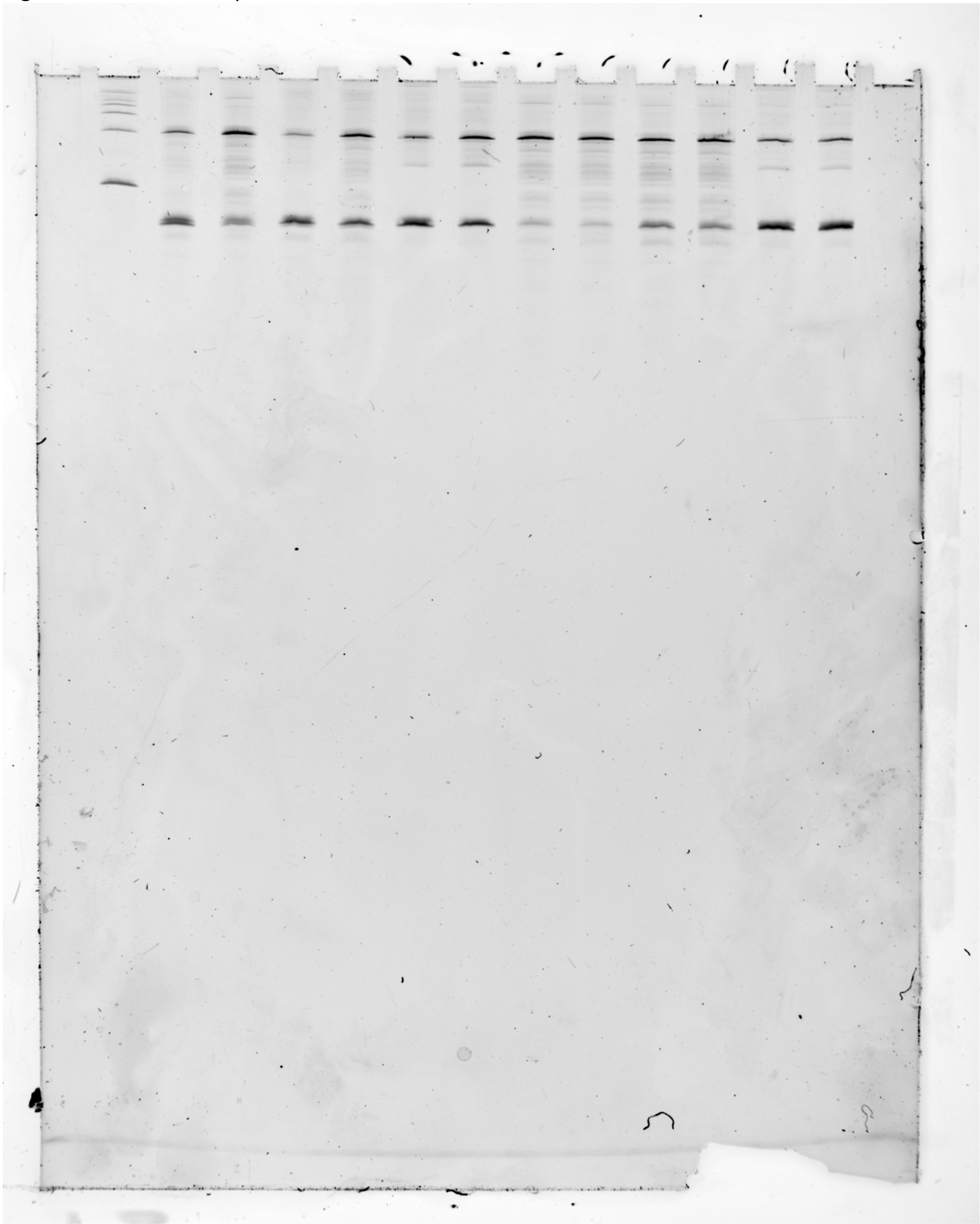

Figure 3E – Notes

3e_Rep1, 3e_Rep2, 3e_Rep3 all show the uncrossed, unprocessed urea-PAGE gel image of the data shown in Fig. 3e, including the other two replicates not explicitly shown in the panel. 3e_Rep3 is the uncrossed, unprocessed image shown in the manuscript. Lanes 1-2 and 15 are not shown in the manuscript.

From left to right:
2. ssRNA Ladder (100, 200, 300, 400, 500, 750, and 1000 bases) (Invitrogen, cat. no. AM7145)
3. In vitro transcript products generated from E. coli RNAp during a single-round of transcription with 1 mM NaCl, DNA template = T. pe Fluoride Riboswitch W1D2
4. In vitro transcript products generated from E. coli RNAp during a single-round of transcription with 1 mM NaF, DNA template = T. pe Fluoride Riboswitch W1D2
5. In vitro transcript products generated from E. coli RNAp during a single-round of transcription with 1 mM NaCl, DNA template = T. pe Fluoride Riboswitch W1D3
6. In vitro transcript products generated from E. coli RNAp during a single-round of transcription with 1 mM NaF, DNA template = T. pe Fluoride Riboswitch W1D3
7. In vitro transcript products generated from E. coli RNAp during a single-round of transcription with 1 mM NaCl, DNA template = T. pe Fluoride Riboswitch W1D4
8. In vitro transcript products generated from E. coli RNAp during a single-round of transcription with 1 mM NaF, DNA template = T. pe Fluoride Riboswitch W1D4
9. In vitro transcript products generated from E. coli RNAp during a single-round of transcription with 1 mM NaCl, DNA template = T. pe Fluoride Riboswitch D2
10. In vitro transcript products generated from E. coli RNAp during a single-round of transcription with 1 mM NaF, DNA template = T. pe Fluoride Riboswitch D2
11. In vitro transcript products generated from E. coli RNAp during a single-round of transcription with 1 mM NaCl, DNA template = T. pe Fluoride Riboswitch D3
12. In vitro transcript products generated from E. coli RNAp during a single-round of transcription with 1 mM NaF, DNA template = T. pe Fluoride Riboswitch D3
13. In vitro transcript products generated from E. coli RNAp during a single-round of transcription with 1 mM NaCl, DNA template = T. pe Fluoride Riboswitch D4
14. In vitro transcript products generated from E. coli RNAp during a single-round of transcription with 1 mM NaF, DNA template = T. pe Fluoride Riboswitch D4

Figure 4 – Raw Gel Replicate 1

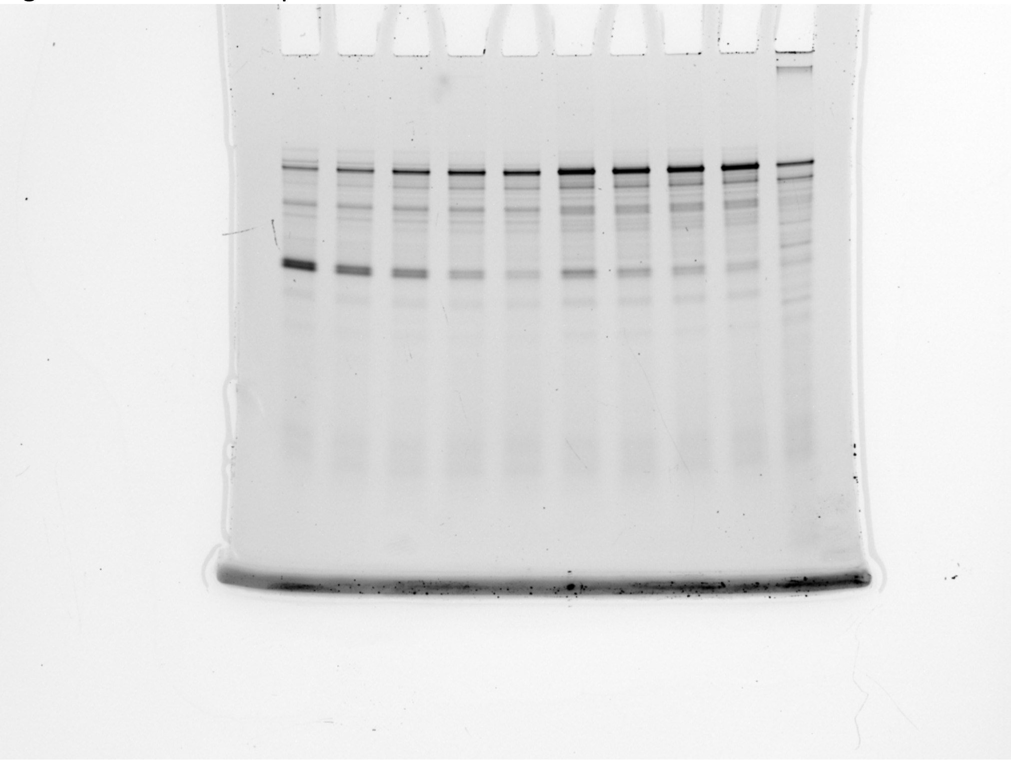

Figure 4 – Raw Gel Replicate 2

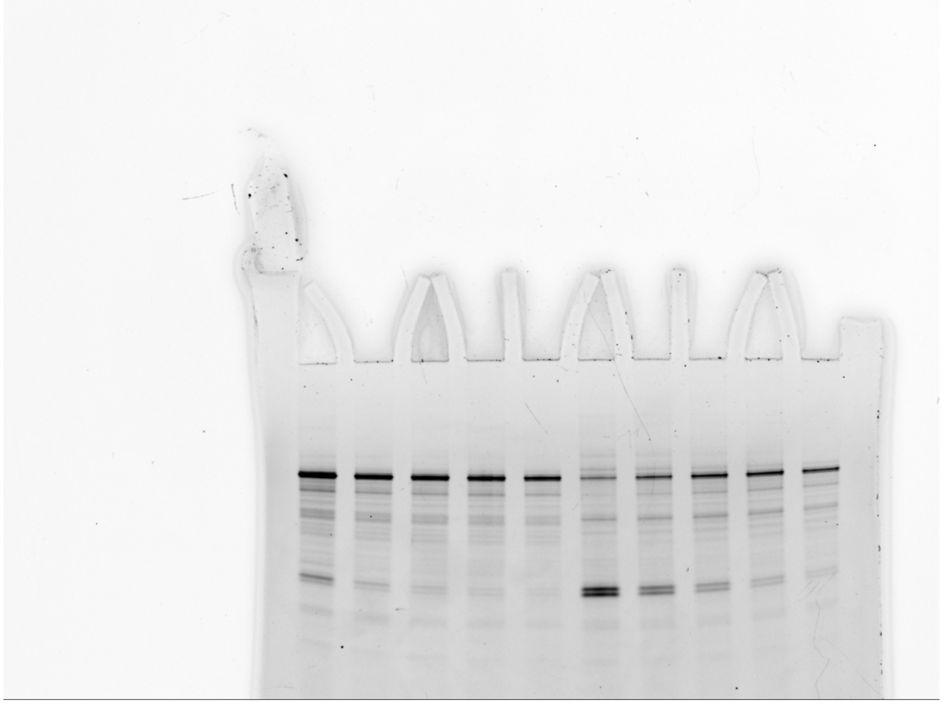

Figure 4 – Raw Gel Replicate 3

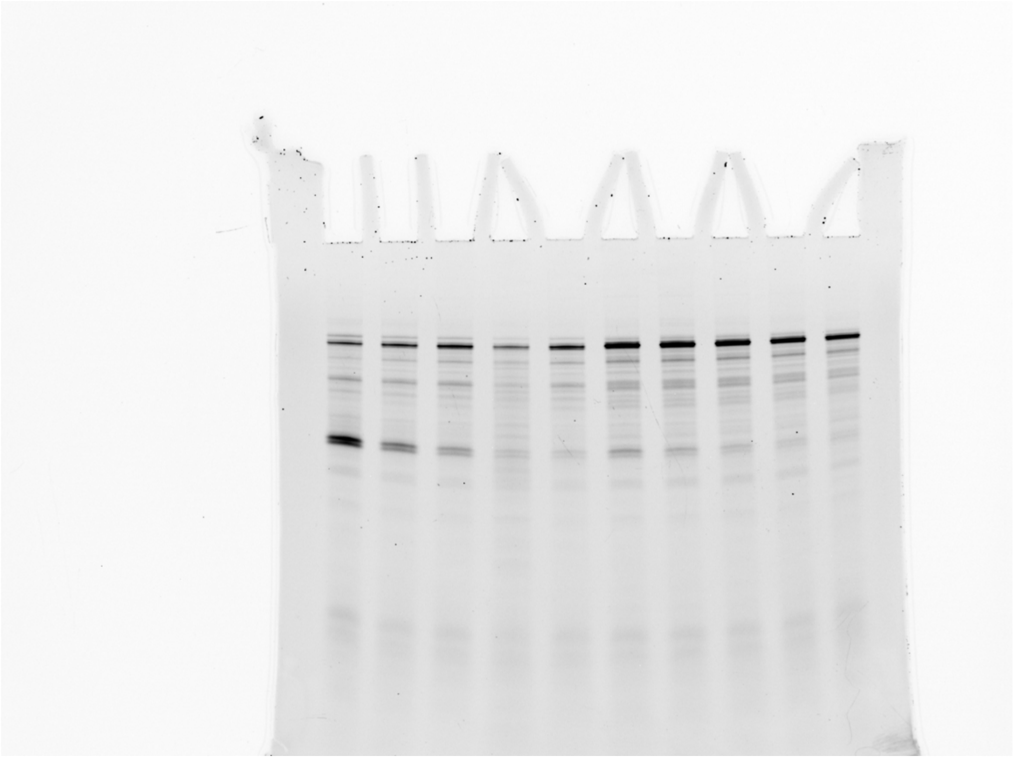

Figure 4 – Raw Gel Replicate 4

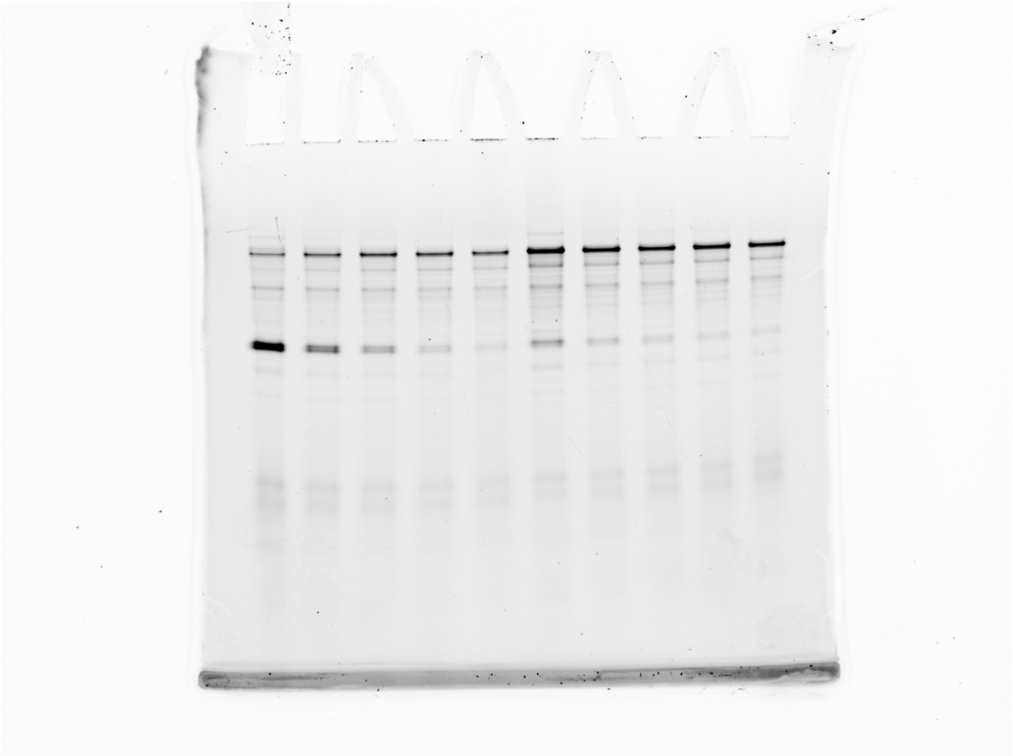

Figure 4 – Notes

4_Rep1, 4_Rep2, 4_Rep3, 4_Rep4 all show the uncrossed, unprocessed urea-PAGE gel image of the data shown in Fig. 4, including the other three replicates not explicitly shown in the panel. 4_Rep4 is the uncrossed, unprocessed image shown in the manuscript.

From left to right (For 4_Rep1, 4_Rep3, 4_Rep4):

1. In vitro transcript products generated by T. aq RNAP in the absence of fluoride, DNA template = T. pe Fluoride Riboswitch W1.
2. In vitro transcript products generated by T. aq RNAP at 1.25 mM NaF, DNA template = T. pe Fluoride Riboswitch W1.
3. In vitro transcript products generated by T. aq RNAP at 2.5 mM NaF, DNA template = T. pe Fluoride Riboswitch W1.
4. In vitro transcript products generated by T. aq RNAP at 5 mM NaF, DNA template = T. pe Fluoride Riboswitch W1.
5. In vitro transcript products generated by T. aq RNAP at 10 mM NaF, DNA template = T. pe Fluoride Riboswitch W1.
6. In vitro transcript products generated by T. aq RNAP in the absence of fluoride, DNA template = T. pe Fluoride Riboswitch WT.
7. In vitro transcript products generated by T. aq RNAP at 1.25 mM NaF, DNA template = T. pe Fluoride Riboswitch WT.
8. In vitro transcript products generated by T. aq RNAP at 2.5 mM NaF, DNA template = T. pe Fluoride Riboswitch WT.
9. In vitro transcript products generated by T. aq RNAP at 5 mM NaF, DNA template = T. pe Fluoride Riboswitch WT.
10. In vitro transcript products generated by T. aq RNAP at 10 mM NaF, DNA template = T. pe Fluoride Riboswitch WT.

From left to right (For 4_Rep2):

1. In vitro transcript products generated by T. aq RNAP in the absence of fluoride, DNA template = T. pe Fluoride Riboswitch WT.
2. In vitro transcript products generated by T. aq RNAP at 1.25 mM NaF, DNA template = T. pe Fluoride Riboswitch WT.
3. In vitro transcript products generated by T. aq RNAP at 2.5 mM NaF, DNA template = T. pe Fluoride Riboswitch WT.
4. In vitro transcript products generated by T. aq RNAP at 5 mM NaF, DNA template = T. pe Fluoride Riboswitch WT.
5. In vitro transcript products generated by T. aq RNAP at 10 mM NaF, DNA template = T. pe Fluoride Riboswitch WT.
6. In vitro transcript products generated by T. aq RNAP in the absence of fluoride, DNA template = T. pe Fluoride Riboswitch W1.
7. In vitro transcript products generated by T. aq RNAP at 1.25 mM NaF, DNA template = T. pe Fluoride Riboswitch W1.
8. In vitro transcript products generated by T. aq RNAP at 2.5 mM NaF, DNA template = T. pe Fluoride Riboswitch W1.

9. In vitro transcript products generated by T. aq RNAP at 5 mM NaF, DNA template = T. pe Fluoride Riboswitch W1.
10. In vitro transcript products generated by T. aq RNAP at 10 mM NaF, DNA template = T. pe Fluoride Riboswitch W1.

Figure S6 – Raw Gel Replicate 1

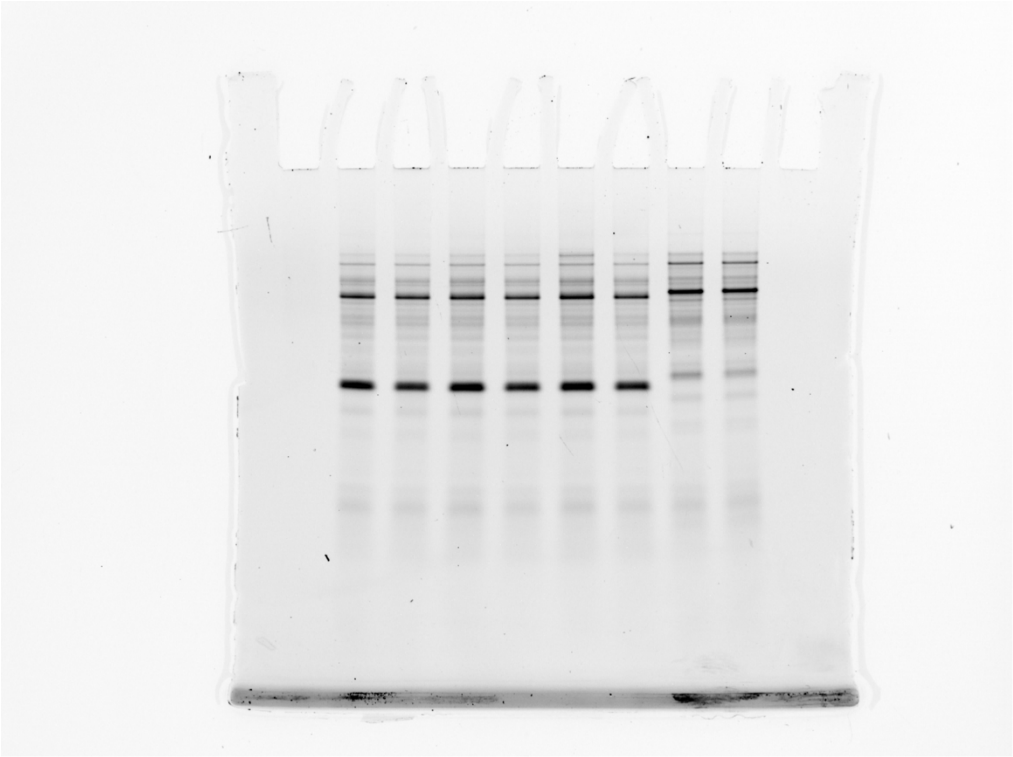

Figure S6 – Raw Gel Replicate 2

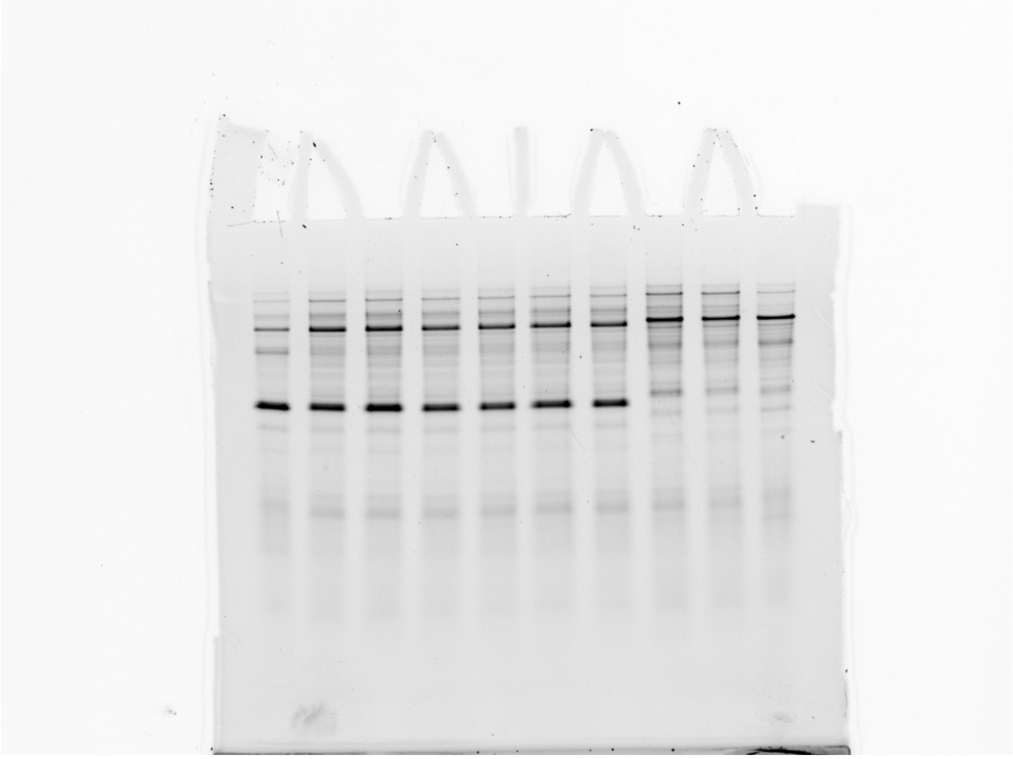

Figure S6 – Notes

S6_Rep1 and S6_Rep2 show the uncrossed, unprocessed urea-PAGE gel image of the data shown in Fig. 4, including the other three replicates not explicitly shown in the panel. S6_Rep1, Lanes 1 and 2, is the uncrossed, unprocessed image shown in the manuscript.

From left to right (For S6_Rep1; Lanes 8-9 are not in the manuscript):
2. In vitro transcript products generated by T. aq RNAP in the absence of fluoride, DNA template = T. pe Fluoride Riboswitch -P1.
3. In vitro transcript products generated by T. aq RNAP at 1 mM NaF, DNA template = T. pe Fluoride Riboswitch -P1.
4. In vitro transcript products generated by T. aq RNAP in the absence of fluoride, DNA template = T. pe Fluoride Riboswitch -P1.
5. In vitro transcript products generated by T. aq RNAP at 1 mM NaF, DNA template = T. pe Fluoride Riboswitch -P1.
6. In vitro transcript products generated by T. aq RNAP in the absence of fluoride, DNA template = T. pe Fluoride Riboswitch -P1.
7. In vitro transcript products generated by T. aq RNAP at 1 mM NaF, DNA template = T. pe Fluoride Riboswitch -P1.

From left to right (For S6_Rep2; Lanes 1, 8-10 are not in the manuscript):
2. In vitro transcript products generated by T. aq RNAP in the absence of fluoride, DNA template = T. pe Fluoride Riboswitch -P1.
3. In vitro transcript products generated by T. aq RNAP in the absence of fluoride, DNA template = T. pe Fluoride Riboswitch -P1.
4. In vitro transcript products generated by T. aq RNAP in the absence of fluoride, DNA template = T. pe Fluoride Riboswitch -P1.
5. In vitro transcript products generated by T. aq RNAP at 1 mM NaF, DNA template = T. pe Fluoride Riboswitch -P1.
6. In vitro transcript products generated by T. aq RNAP at 1 mM NaF, DNA template = T. pe Fluoride Riboswitch -P1.
7. In vitro transcript products generated by T. aq RNAP at 1 mM NaF, DNA template = T. pe Fluoride Riboswitch -P1.
